# Sugar ABC transporter repertoires predict ecological dynamics in gut microbiome communities

**DOI:** 10.1101/2025.11.12.686590

**Authors:** Harsh Maan, William Jogia, Caichen Duan, Fanny Matheis, Eric K. Nishimoto, Chenzhen Zhang, Alexis P. Sullivan, Jonas Schluter

## Abstract

The gut microbiome plays a central role in human health, but modern diets and lifestyles alter its composition. The microbial genomic traits that drive these ecological shifts, particularly in response to dietary sugars, remain poorly characterized. Here, we integrate a large dataset of longitudinal human diet-microbiome records and comparative genomics of human and murine gut isolates with *in vitro* and *in vivo* experiments to identify sugar ABC (ATP-binding cassette) transporters as key predictors of bacterial fitness and microbial community responses to dietary sugars. Strains encoding these transporters exhibit enhanced growth and consistently outcompete others in both monocultures and complex consortia across contexts. In gnotobiotic mice, dietary sugar supplementation selectively increases the expansion of sugar ABC transporter-positive bacteria, including the model gut pathobiont *Escherichia coli*. Systematic deletion of sugar transporter genes in *E. coli* revealed that a specific sugar ABC transporter gene was required to invade a model gut consortium, highlighting its importance in microbial competition. Together, these findings establish sugar ABC transporters as genomic predictors of microbial community dynamics in response to dietary sugars.

## Introduction

Gut microbiome composition is associated with regulation of metabolism, immunity, and overall health^1,2^. Modern lifestyle factors, such as environmental changes, widespread antibiotic use, and changes in diet, including an increase in sugar consumption, are disrupting the human gut microbiome symbiosis, causing lasting multi-generational shifts in microbial composition^1,3,4^. Altered microbiome compositions are associated with increasing rates of chronic diseases, including colitis, metabolic syndrome, obesity, type 2 diabetes, and inflammatory bowel disease^4–6^. Emerging evidence suggests that Western diets high in sugars, fats and low in fiber can reduce microbial diversity and alter bacterial gene repertoires^7^, and dietary sugars can also directly silence gene expression of gut colonization factors in bacterial symbionts^8^. Despite the growing evidence linking sugar consumption to microbiome disruption, the genetic and ecological mechanisms underpinning bacterial responses to dietary sugars remain largely unexplored.

Predicting complex microbial ecosystem responses to changing diets is challenging as they may be nonlinear, context-dependent, and hysteretic, meaning they depend on prior ecological interactions and environmental conditions^9,10^. Moreover, while advances in sequencing and computational modeling have identified correlations between high-sugar diets and microbiome shifts, the molecular mechanisms dictating which microbes thrive or decline have yet to be fully defined. In murine models, high sugar intake increased Pseudomonadota abundance while decreasing Bacteroidota^11^, liquid fructose-rich elevated the Bacillota-to-Bacteroidota ratio^12^, whereas a high-fat, high-sucrose diet reduced Bacteroidota and increased Bacillota abundance, which partially depended on host genetic background^13^. Human studies comparing rural African children who consumed high-fiber diets with European children on sugar-rich Western diets showed that fiber increased Bacteroidota diversity, while Western diets raised Bacillota abundance and lowered overall microbiome diversity^14^. Clinically, patients undergoing hematopoietic cell transplantation who consumed high-sugar diets experienced exacerbated antibiotic-induced loss of microbial diversity and expansion of antibiotic-resistant *Enterococcus* species^3^. While these studies reveal strong associations between sugar intake and microbiome changes, the altered microbial components vary from study to study, and the genetic determinants driving microbial fitness in sugar-rich diets—which could therefore consolidate these observations—remain elusive.

A critical driver of microbial growth is the ability to access and metabolize sugars^15,16^. To import sugars efficiently, bacteria employ a range of transporters including ATP-binding cassette (ABC) transporters, phosphotransferase system (PTS) transporters and Major Facilitator Superfamily (MFS) transporters^17^, which enable entry into key glycolytic pathways like the Embden-Meyerhof-Parnas (EMP) pathway, the Entner-Doudoroff (ED) pathway, and the pentose phosphate pathway (PPP)^18^. The contribution of each transporter family in sugar uptake has been characterized in individual species^17^, revealing that their activity is often species-specific and influenced by environmental conditions. For instance, in some strains of *E. coli*, sugar ABC transporters are highly expressed under low-glucose conditions, while PTS transporters dominate when glucose was abundant^19–21^. In contrast, *in vitro* studies on *Enterococcus faecium* indicate it depends on PTS transporters for growth and biofilm formation^22^. In uropathogenic *E. coli*, ABC transporters deletion reduced *in vitro* growth rates and reduced fitness in infection models^23^. In *Pseudomonas stutzeri*, ABC transporters for glucose uptake were crucial for growth in both low and high sugar environments^24^. These results suggest a key role of transporter systems in microbial competition, which we hypothesize is crucial in the complex and dynamic gut environment.

In this study, we combined genomic analyses of human and mouse gut bacteria with *in vitro* and *in vivo* experiments to demonstrate that sugar ABC transporters are determinants of bacterial competitiveness in model microbial communities. To assess whether these findings extend to the human microbiome, we analyzed paired microbiome and dietary intake data across 9,419 meals and 1,009 stool samples from hematopoietic cell transplant (HCT) patients and found that the abundance of bacterial genera harboring sugar ABC transporters correlates with the amount of dietary sugar intake. We colonized gnotobiotic mice with a defined gut consortium, and observed that strains encoding sugar ABC transporters expanded significantly when we provided the mice with sugar-supplemented water, akin to sugary drinks. Finally, using targeted gene deletions in *E. coli*, we show that deletion of a single ABC transporter gene significantly impaired invasion of a resident community. These findings establish an outsized importance of sugar ABC transporters as predictive genomic markers for microbial ecosystem responses to dietary sugars, and that they are critical for ecological success. Together, our work is advancing genetics-driven frameworks towards mechanistic forecasting of microbiome dynamics.

## Results

### Interphylum genomic characterization of sugar transporter repertoires and metabolic pathways in gut bacteria

As core metabolic pathways in bacteria are highly conserved^25,26^ but nutrient transporter systems vary even between related strains^23^, we hypothesized that sugar transporters and the core metabolic pathways they support would be under separate selection, which would decouple their distributions across bacterial genomes. To test this, we analyzed 1,147 bacterial genomes from five major phyla, broadly representing gut bacterial diversity^1^ (Supplementary Fig. **1**). Our analysis focused on six core genomic features involved in the utilization of simple sugars: three sugar transport systems: ATP binding cassette (ABC), Major Facilitator Superfamily (MFS), and phosphotransferase system (PTS); and three central carbon metabolic pathways: Embden-Meyerhof-Parnas (EMP), pentose phosphate pathway (PPP), and Entner-Doudoroff (ED) pathways) (Fig. **1a**). Visualization of gene presence on a tree constructed from the analyzed genomes and pairwise mutual information (PMI) analysis of gene presence across phyla revealed only partial overlap between transporter and metabolic pathway modules (Fig. **1a**). Among species of the same phylum, we observed considerable variation in transporter and glycolytic pathway presence (except Verrucomicrobiota, represented solely by *Akkermansia muciniphila*; Extended Data Fig. **1**), indicating that sugar metabolism traits are not strictly conserved phylogenetically.

**Figure 1.**
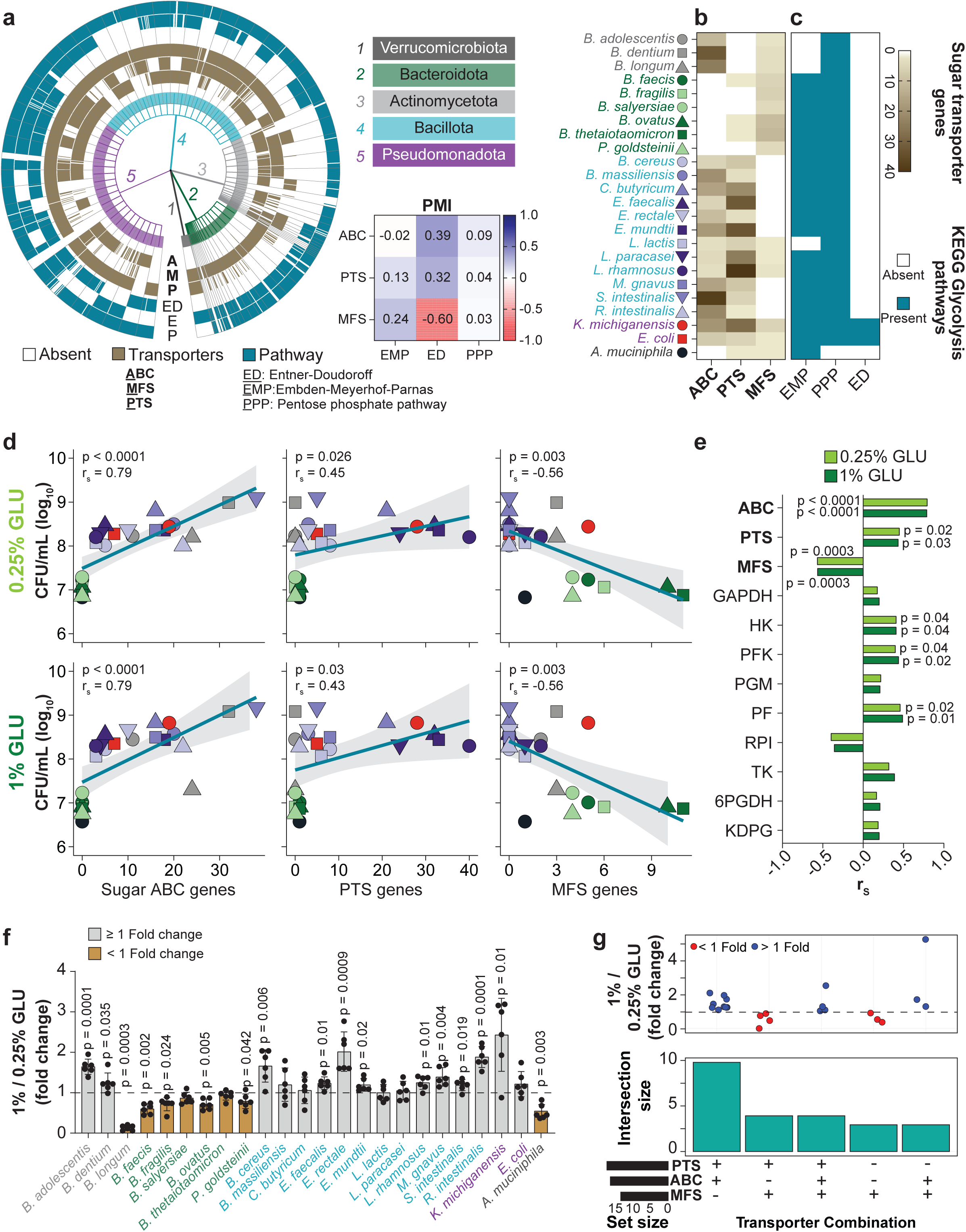
Independent variation in sugar utilization and transporter gene repertoires explain growth phenotypes of bacteria across phyla. **a)** Sugar transporter (PTS, MFS, ABC) and glycolytic pathways (PPP, EMP, ED) vary independently within and across phyla. The table reports pairwise mutual information (PMI) values between indicated features, analyzed from 1,147 gut bacterial genomes visualized on the tree graph. **b)** Sugar transporter gene counts in genomes of 24 representative species. **c)** Presence of complete glycolytic pathways in the genomes of the 24 species, as determined by KEGG pathway reconstruction. **d)** Spearman correlation (rₛ) between colony-forming units (CFUs) and sugar transporter gene counts for 24 strains grown in BHI-NG medium supplemented with 0.25% or 1% glucose (GLU), harvested at 24 h post inoculation. Initial inoculum at 0 h was 10^5^ CFU/mL ± 0.3. *P*-values: Spearman correlation; lines: linear regression (Ordinary Least Squares, OLS); shaded areas: 95% CIs. **e)** Spearman correlations (rₛ) between gene counts and CFUs for sugar transporter genes and glycolytic pathway genes at 24 h growth. **f)** Mean ± SD fold change in CFUs between 0.25% and 1% GLU at 24 h for each species. Brown bars highlight species exhibiting a fold change < 1. *P*-values: two-tailed one-sample ratio t-test against a fold change of 1. **g)** UpSet plot of transporter repertoires in the 24 gut bacterial species. Dot plots depict mean fold changes in growth upon GLU increase from 0.25% to 1%. Fold changes for species corresponding to brown bars in panel f were now obtained using a pH-adjusted 1% GLU condition (pH 6.8), while fold changes for grey bars are identical to those shown in panel f. The horizontal line at fold change = 1 denotes no difference in growth between glucose concentrations. Set size indicates the total number of species carrying each transporter, and intersection size indicates the number of species sharing the corresponding combination of transporters. For panels **d–g**, data are combined from two independent biological experiments with three technical replicates each (total n = 6). *P* < 0.05 was considered statistically significant.

To evaluate how this genomic variability translates to functional differences, we selected 24 genetically diverse gut bacterial species from the five major phyla for experimental testing (Supplementary Table **1**). Among these strains, sugar transporter profiles showed both phylum-level patterns and intra-phylum variability (Fig. 1**b**, Supplementary Fig. **2**), reflecting the full genome data set (Fig. 1**a**). Pseudomonadota encoded comparable numbers of sugar ABC and PTS transporters. Bacillota showed heterogeneity, with sugar ABC and PTS transporters variably present across species, while MFS transporters were largely absent. Bacteroidota primarily encoded MFS transporters, with some strains also possessing PTS transporters, and consistently lacked sugar ABC transporters. Actinomycetota possessed both sugar ABC and MFS transporters but lacked PTS systems, whereas *Akkermansia muciniphila* (Verrucomicrobiota) encoded only PTS and MFS transporters. Glycolytic metabolism repertoires also varied by phylum and species (Fig. 1**c**). The EMP pathway was nearly ubiquitous, absent only in *Lactococcus lactis* (Bacillota) and Actinomycetota species. The PPP pathway was broadly present but absent in *A. muciniphila*. The ED pathway only occurred in Pseudomonadota. To evaluate functional completeness, three essential enzymes were examined for each pathway (Supplementary Table **2**). No genome encoded the full set of representative enzymes for all three pathways. Gene presence patterns varied across species, with some pathways missing specific enzymes and others sharing enzymes between pathways, reflecting the modular and overlapping organization of glycolytic pathways^27^ (Supplementary Fig. **3**). This heterogeneity in sugar metabolism genes and transporters did not align with phylogeny, reinforcing that sugar utilization traits are at least partially decoupled from taxonomy and may have been shaped by strain-specific selective pressures. Taken together, our selected strains encode independent variation in sugar metabolism and transport systems.

### Growth in simple sugars correlates with sugar ABC transporter repertoires

With this, we next sought to determine which of these genomic features correlate with growth on a permissive medium with controlled simple sugar concentrations. We cultured each of the 24 bacterial species individually in a modified Brain Heart Infusion (BHI) medium lacking glucose (BHI-NG), and supplemented it with either 0.25% or 1% (w/v) glucose (GLU), reflecting the nutrient variability in gut environments^4,28^. After 24 hours of growth, colony-forming units (CFUs) were quantified and correlated with genes involved in sugar transport and processing (Fig. **1d**). Growth correlated most strongly with sugar ABC transporter gene counts (Spearman’s *r_s_* = 0.79, *p* < 0.0001), while MFS transporter gene counts were negatively correlated with growth (*r_s_* = –0.56, *p* = 0.0003), with weaker associations between CFUs and central metabolism gene counts; our results were consistent in both glucose concentrations (Fig. **1d**, **e** and Extended Data Fig. **2**). Next, to assess if increased sugar availability altered growth in a genetic feature-specific manner, we compared 24 h CFU ratios between 1% and 0.25% GLU. We found that some species exhibited reduced growth in 1% GLU (Fig. **1f**). As even minor differences in pH may selectively inhibit strains^29,30^, we adjusted the pH of 1% GLU medium from 6.30 to 6.80 (matching 0.25% GLU), which significantly rescued *B. longum* growth (*p* = 0.0008, Supplementary Fig. **4**), moderately improved *A. muciniphila*, which had shown a significant growth reduction (Fig. **1f**), but failed to significantly improve growth in Bacteroidota (Supplementary Fig. **4**). Analysis of transporter combinations after pH adjustments revealed that bacteria harboring sugar ABC transporters consistently benefitted from higher glucose concentration (Fig. 1**g**).

This observation was also robust to a binary classification of the strains by presence or absence of transporter types; strains encoding sugar ABC transporters exhibited significantly higher growth in both low and high glucose conditions compared to those lacking them (Extended data Fig. **3** and **4**). Moreover, multivariable analysis of CFU counts with gene presence/absence at 12, 24, and 48 h showed that presence of ABC sugar transporters had the strongest and most significant association with CFU among all genetic features, except at 48 h in 0.25% GLU (Fig. **2a**, Supplementary Fig. **5**). We extended our experiments to other simple dietary sugars (Fig. **2b**, Supplementary Fig. **6****-9**); using area under the curve (AUC) as a reliable proxy for bacterial growth (Fig. **2c**), we found that species with sugar ABC transporters grew better on fructose (FRU), sucrose (SUC), and galactose (GAL).

**Figure 2.**
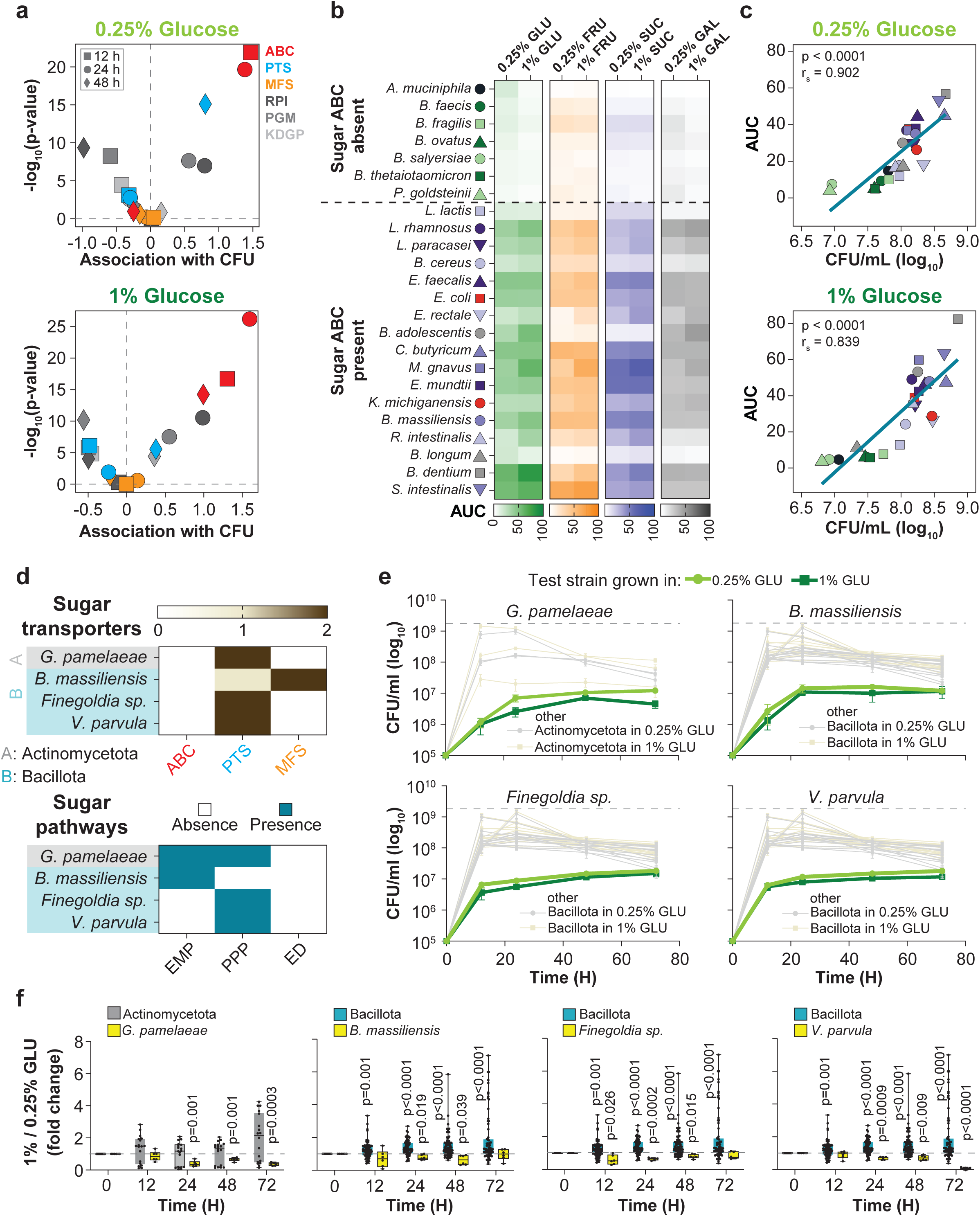
Bacteria with sugar ABC transporters exhibit enhanced *in vitro* growth on simple sugars. **a)** Effect sizes and statistical significance for gene presence predictors of CFU counts in 0.25% and 1% GLU, estimated using a multivariable linear mixed-effects model with gene presence repertoires as fixed effects and a varying intercept for technical repeats. **b)** Area under the curve (AUC) from planktonic growth (OD_600_) of indicated species cultured in BHI-NG medium supplemented with 0.25% or 1% GLU, fructose (FRU), sucrose (SUC), or galactose (GAL) over 72 h. **c)** Spearman correlation analysis examining the relationship between AUC and average CFUs measured over 72 h. **d)** Distribution of genes involved in sugar transport and presence or absence of glycolytic pathways in the indicated bacteria. **e)** Growth curves of indicated species lacking sugar ABC transporters compared with growth of other bacteria within the same phylum (faint lines) that possess sugar ABC transporters (see also Supplementary Fig. 5); dashed lines represent the maximum growth achieved by members of the respective phylum under identical conditions. **f)** Fold-differences of CFUs (1% / 0.25% GLU) for indicated species (yellow boxes) and their phylum counterparts (grey boxes) across different time points. Each group was independently compared against a fold change of 1 using a two-tailed one-sample ratio t-test. For panels **a–e**, data are combined from two independent biological experiments with three technical replicates each (total n = 6). *P* < 0.05 was considered statistically significant.

As sugar ABC transporters seemed to accelerate growth, we next tested strains that, unlike other members of their phylum in our species set, lacked sugar ABC transporters, expecting slower growth compared to their relatives from the same phylum (Fig. **2d**). The new strains encoded genes for PTS and MFS transporters and at least one complete glycolytic pathway, but, as predicted, grew worse than their sugar ABC transporter possessing counterparts from the same phylum (Fig. **2e**, Extended data Fig. **5**). Moreover, these ABC-lacking strains did not benefit from elevated glucose concentrations (Fig. **2f**), in contrast to their phylum-matched counterparts. Together, these results support a model whereby transporter repertoire, rather than taxonomy, explains growth on simple sugars.

### Taxa associated with dietary sugar intake in HCT patients harbor abundant sugar ABC transporters

We thus hypothesized that the repertoire of sugar ABC transporter genes could serve as a useful marker for predicting ecological patterns in complex microbial communities responding to dietary sugars. To investigate this, we analyzed human microbiome data from fecal samples of patients undergoing HCT, in a cohort with high-resolution dietary intake data^3^. This data set comprises per meal macronutrient intake, including sugars, as well as food-item level resolution revealing the frequent consumption of sugar-sweetened beverages (Supplementary Table **3**).

For each microbiome sample, we modeled the association between total sugar intake in the prior two days^3^ and the centered log ratio (CLR)-transformed relative abundances of 91 bacterial genera using Bayesian regression models (Extended data Fig. **6**; n = 1,009 samples from 158 patients). As this amplicon-based microbiome data set does not directly reveal transporter gene abundances, we estimated them by computing the average count of each sugar transporter type from up to 100 randomly sampled complete genomes per genus from the NCBI genome database. This revealed a consistent pattern: genera with confidently positive sugar intake associations exhibited a higher total number of sugar transporter genes compared to genera negatively associated with sugar intake (Fig. **3a**). The largest positive coefficient was obtained for the genus *Blautia,* with an average computed sugar ABC transporter count of 46 genes. Conversely, of typical gut commensals, we found *Prevotella*^2,31,32^ to be most negatively associated with dietary sugar intake, and it lacks sugar ABC transporters (Fig. **3b**). This suggested that sugar ABC transporters were specifically predictive of genus abundance associations with dietary sugar consumption. Thus, we next correlated the average transporter count per genus of all sugar transporter genes combined, as well as each transporter type individually, with the dietary sugar association coefficients for the corresponding genera. Both Spearman rank correlations and Bayesian linear regressions showed that total sugar transporter counts and specifically the sugar ABC transporter counts (Fig. **3c**) were significantly and positively correlated with the dietary sugar association coefficients. In contrast, PTS and MFS sugar transporter counts did not show this relationship (Fig. **3c**). This analysis validated the predictive power of genomic sugar ABC transporter counts, which emerged as the strongest predictor of diet-abundance associations in human microbiomes responding to dietary sugars.

**Figure 3.**
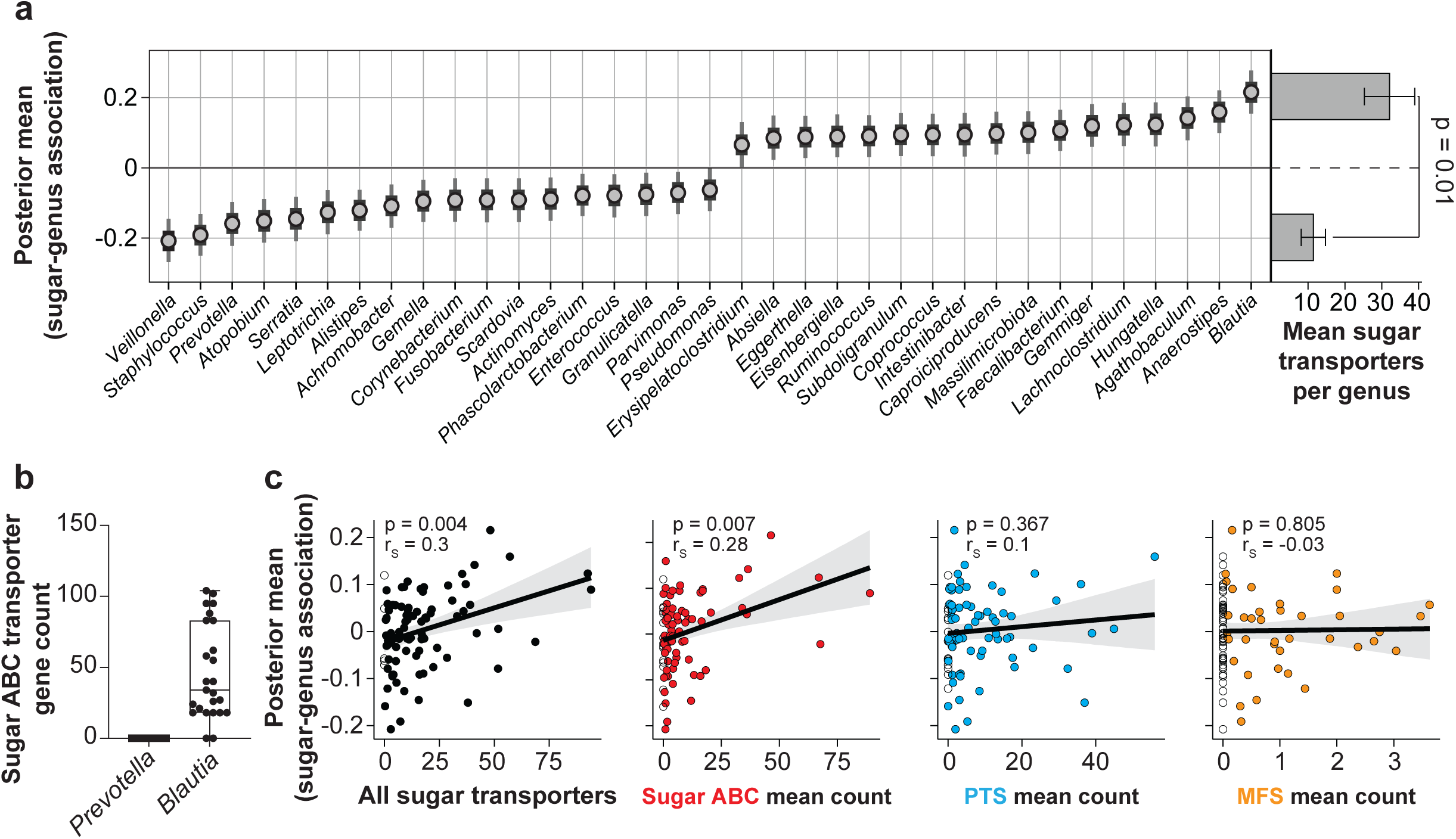
In the human gut microbiome, sugar ABC transporter repertoire is associated with response to dietary sugar intake. **a)** Bayesian linear regression posterior coefficient distributions of genus associations with sugar intake recorded during the preceding 2 days in patients undergoing hematopoietic cell transplantation (HCT) (n = 1,009 samples from N = 158 patients^3^, lines: 66% and bars: 95% posterior Credible Intervals, C.I.). Only genera for which the 95% highest posterior density interval did not include zero are shown (35 of 91 genera; full results in Extended Data Fig. 6). Right, bar plots showing the mean number of sugar transporter genes from randomly sampled complete genomes in NCBI for genera positively (top, n = 17) and negatively (bottom, n = 18) associated with sugar intake. Statistical comparison was performed using a two-sided Wilcoxon rank-sum test. **b)** Sugar ABC transporter gene counts across genomes of the genera *Blautia* (n=25) and *Prevotella* (n = 66), two common gut symbionts. **c)** Bayesian linear regressions of average sugar transporter gene counts per transporter type and genus–sugar association coefficients for all genera with ≥1 complete NCBI genome (n = 91, as shown in panel a and Extended Data Fig. 22). Lines and shaded area: posterior means with 95% C.I.. Spearman correlation coefficients are reported in the upper left. Open circles indicate genera with zero transporters of the corresponding type.

### Sugar ABC transporters are associated with growth in *in vitro* communities

Building on monoculture results and observational human data, we hypothesized that sugar ABC transporters drive ecological success in microbial communities under sugar supplementation. To test this, we leveraged a 15 species Oligo-Mouse-Microbiota (OMM^15^), a minimal consortium representing five dominant murine gut phyla^33,34^, whose genomes varied in transporter repertoires (Supplementary Fig. **10**, Supplementary Table **4**). To ensure comparability to results from the human strains, we assessed monoculture strain growth of this new library of murine strains in our growth assays. Two strains (*Acutalibacter muris* and *Streptococcus danieliae*) did not grow on BHI-NG because they require blood-based and Tryptic Soy Broth media, respectively^35^. Among the other strains, consistent with our previous experiments, strains harboring sugar ABC transporters showed enhanced monoculture growth across GLU concentrations (Fig. **4a**) and other simple sugars (Fig. **4b**, Supplementary Figs. **11****-13**).

**Figure 4.**
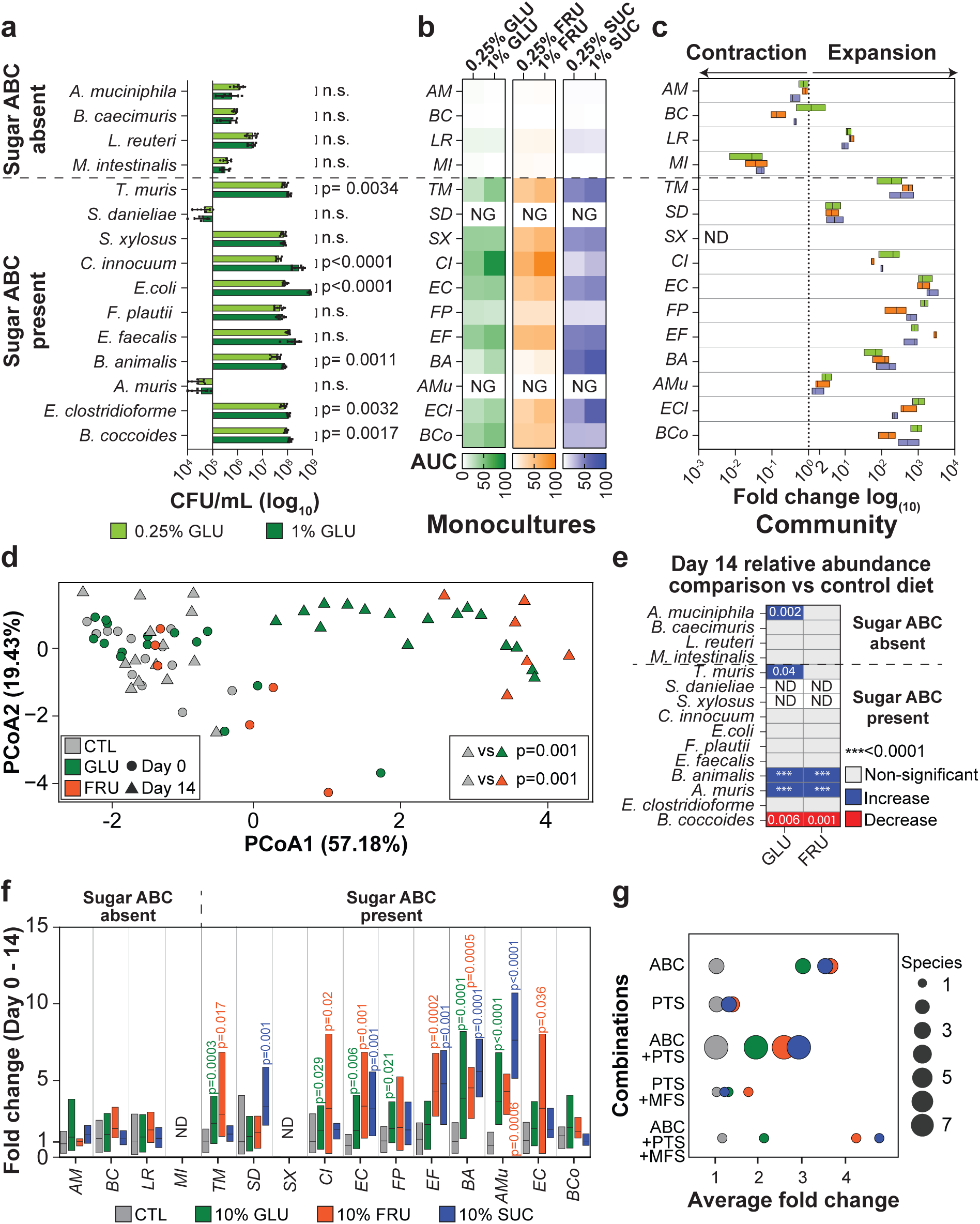
Sugar ABC transporters predict microbial community dynamics of OMM^15^ strains *in vitro* and in gnotobiotic mice. **a)** Mean ± SD CFUs of the indicated species after 48 h. Statistical comparison was performed between 0.25% and 1% GLU for each species using an unpaired t-test with Welch’s correction. **b)** Area under the curve (AUC) from planktonic growth (OD_600_) of the indicated species cultured in BHI-NG medium with 0.25% or 1% GLU, FRU or SUC over 72 h. NG, no planktonic growth. **c)** Fold change in density of OMM^15^ strains grown *in vitro* from 0 to 72 h, in BHI-NG supplemented with 0.25% GLU, FRU and SUC, measured by quantitative PCR (qPCR). Fold change >1 after 72 h indicates expansion. ND, not detected. For panels **a** and **b**, data are combined from two independent biological experiments with three technical replicates each (total n = 6). For panel **c,** data are combined from two independent biological experiments and qPCR was performed in technical duplicate for each species (n = 4). **d)** Principal Coordinates Analysis (PCoA, Aitchison distance) of fecal microbial community composition from gnotobiotic mice based on relative abundances of taxa in control, 10% GLU and FRU treatment groups at baseline (day 0) and day 14. PERMANOVA analysis at day 14 revealed significant compositional differences between treatments: control vs. GLU (p = 0.001) and control vs. FRU (p = 0.001). **e)** Differences of OMM15 species’ relative abundances on day 14 between mice on a sugar-supplemented or control diet. Directionality of differences with adjusted p-values < 0.05 is indicated by color; Kruskal–Wallis test with Dunn’s multiple comparisons. ND: not detected. For panels **d and e**, data are combined from two independent experiments (n = 8 control female mice, n = 7 control male mice, n = 7 GLU female mice, n = 9 GLU male mice, n = 3 FRU female mice, n = 3 FRU male mice). **f)** Fold changes in density of OMM^15^ species from day 0 to day 14, calculated from normalized qPCR Ct values (Ct/fecal weight) and stratified by diet groups. Each bar represents the distribution of the observed within-mouse fold change for the indicated species from day 0 to day 14. Statistical analysis comparing control vs sugar-supplemented density changes was performed using the Kruskal–Wallis test with Dunn’s multiple comparisons, with adjusted *p*-values shown. **g)** Average fold change in species density from day 0 to day 14 grouped by transporter combinations. Circle color denotes treatment condition; circle size indicates the number of species sharing the corresponding transporter profile. For panels **f** and **g**, data are combined from three independent experiments (n = 10 control female mice, n = 9 control male mice, n = 7 GLU female mice, n = 9 GLU male mice, n = 3 FRU female mice, n = 3 FRU male mice, n = 3 SUC female mice, n = 2 SUC male mice). *M. intestinale* and *S. xylosus* were below detection by qPCR in fecal samples; For panels **f** and **g**, qPCR was performed in technical duplicate for each species. ND, not detected.

Before testing this consortium *in vivo* we sought to assess how community context altered each strains growth on different sugars *in vitro.* We inoculated approximately 10^3^ ± 0.3 CFUs/ml of each of the 15 strains simultaneously into BHI-NG medium supplemented with either 0.25% GLU, FRU, or SUC. We monitored species’ density changes over a 72-hour period using quantitative PCR (qPCR) with primers specific to each strain’s 16S rRNA gene, defining a fold change from 0 to 72 h of greater than 1 as growth within the community (Fig. **4c**). All sugar ABC transporter carriers except for *S. xylosus* expanded: most notably, *E. coli* and *Enterococcus faecalis* grew rapidly during the earlier time points (8 h) with other following around 12 h (Supplementary Fig. **14**). In contrast, bacteria lacking sugar ABC transporters did not expand in the community despite their ability to expand in monocultures (Fig. 4**a**). An exception was *Limosilactobacillus reuteri,* which possesses PTS transporters and showed modest growth in the community settings. *A. muris* and *S. danieliae*, which failed to grow in monoculture, showed weak expansion in the community (Fig. **4c** and Supplementary Fig. **14**). Taken together, sugar ABC transporters were predictive of growth in different simple sugar conditions in a defined *in vitro* community of diverse murine gut bacterial strains.

### Sugar ABC transporters are predictive of sugar-induced expansion *in vivo*

We hypothesized that sugar ABC transporter profiles could forecast microbiome dynamics under sugar enriched diets. To test this, we colonized germ-free mice with the OMM^15^ community and supplemented their drinking water with 10% GLU, FRU, or SUC for 14 days, modeling typical sugar concentrations in hospital or commercial beverages^5,36^. Both the control and sugar-fed groups were maintained under otherwise identical conditions in the same room, housed in HEPA-filtered ISOcages in a gnotobiotic facility, with *ad libitum* access to their drinking water. As reported previously, sugar-fed mice consumed more water (data not shown), gained weight, and exhibited reduced colon length and cecal weight compared to controls (Supplementary Figs. **15****-17**)^5,37^. To assess the impact of dietary sugars on the gut microbiota, we collected fecal samples at days 0 and 14 and profiled community compositions with 16S rRNA gene sequencing. Principal coordinates analysis (PCoA) showed that by day 14, the microbiota of GLU- and FRU-treated mice had diverged significantly from that of control mice as well as from their own baseline microbiota (Fig. **4d** and Supplementary Fig. **18**). Analysis of relative abundances at day 14 between sugar treated mice and control revealed a significant relative abundance increase of *A. muris*, *Bifidobacterium animalis*, *Turicibacter muris*, and *A. muciniphila* in sugar-treated groups, while *Blautia coccoides* showed a significant decrease (Fig. **4e**). These findings demonstrate that dietary sugar intake drives substantial restructuring of this model consortium *in vivo*. A known limitation of 16S-based community profiling is that it provides compositional data, obscuring the biological interpretation of observed changes due to negative covariance biases^38,39^. To address this, we implemented complementary qPCR to directly measure changes in density of bacterial strain between days 0 and 14, providing a direct measurement of population dynamics following sugar supplementation. This revealed significant differences between control and sugar treated mice (Supplementary Fig. **19**), including increases of sugar ABC transporter-positive *B. animalis* and *A. muris* in all sugar-supplemented diets. Consistent with this, individual strain dynamics from day 0 to day 14 within each mouse (Fig. **4f**) revealed significantly greater expansion of most sugar ABC transporter-positive strains in sugar-fed mice relative to mice on control diets, but none of the strains without sugar ABC transporters showed sugar-induced increased growth. Combining strains based on their transporter combinations confirmed that bacteria with sugar ABC transporters, exclusively or with other transporters, expanded more *in vivo* upon sugar supplementation (Fig. **4g**, Extended data Fig. **7**). Thus, we hypothesized that sugar ABC transporters are crucial drivers of bacterial competitiveness under sugar supplementation.

### A specific sugar ABC transporter is required for community invasion by *E. coli*

Microbiome invasion by pathobiontic proteobacteria is a pattern of dysbiosis^40,41^, and resource competition can limit their expansion^34,42^. To explored the necessity of specific transporters for invasion of a resident gut community, we created deletion mutants (Supplementary Table **5**) for each transporter gene of the proteobacterial OMM^15^ strain, *E. coli MT1B1* and compared their growth in monocultures to wild-type (WT). In 0.25% glucose, deletions of *mglB* (a sugar ABC transporter subunit) and *ptsG1* (a PTS transporter subunit) resulted in the strongest growth defects (Fig. **5a**, Supplementary Fig. **20**). To compare mutant fitness in a community, we separately inoculated WT *E. coli* and its Δ*mglB* and Δ*ptsG1* mutants together with the other OMM^14^ community members. While both Δ*ptsG1* and Δ*mglB* mutants showed reduced expansion relative to WT initially, the Δ*ptsG1* mutant eventually caught up, showing no significant expansion differences to the WT after 24 hours (Fig. **5b**), in contrast to Δ*mglB*, which expanded less than both WT and Δ*ptsG1.* We next assayed community invasion fitness by first establishing a OMM^14^ community lacking *E. coli* and introducing the *E. coli* WT or the mutant strains after 12h, i.e. when glucose concentrations reached ≈50% of the initial concentrations (Fig. **5c**, Supplementary Fig. **21**). The Δ*mglB* mutant showed significantly lower competitiveness in this assay than both WT and Δ*ptsG1* strains at 36 hours and 60 hours (Fig. **5d**), highlighting the importance of a single sugar ABC transporter for community invasion.

**Figure 5.**
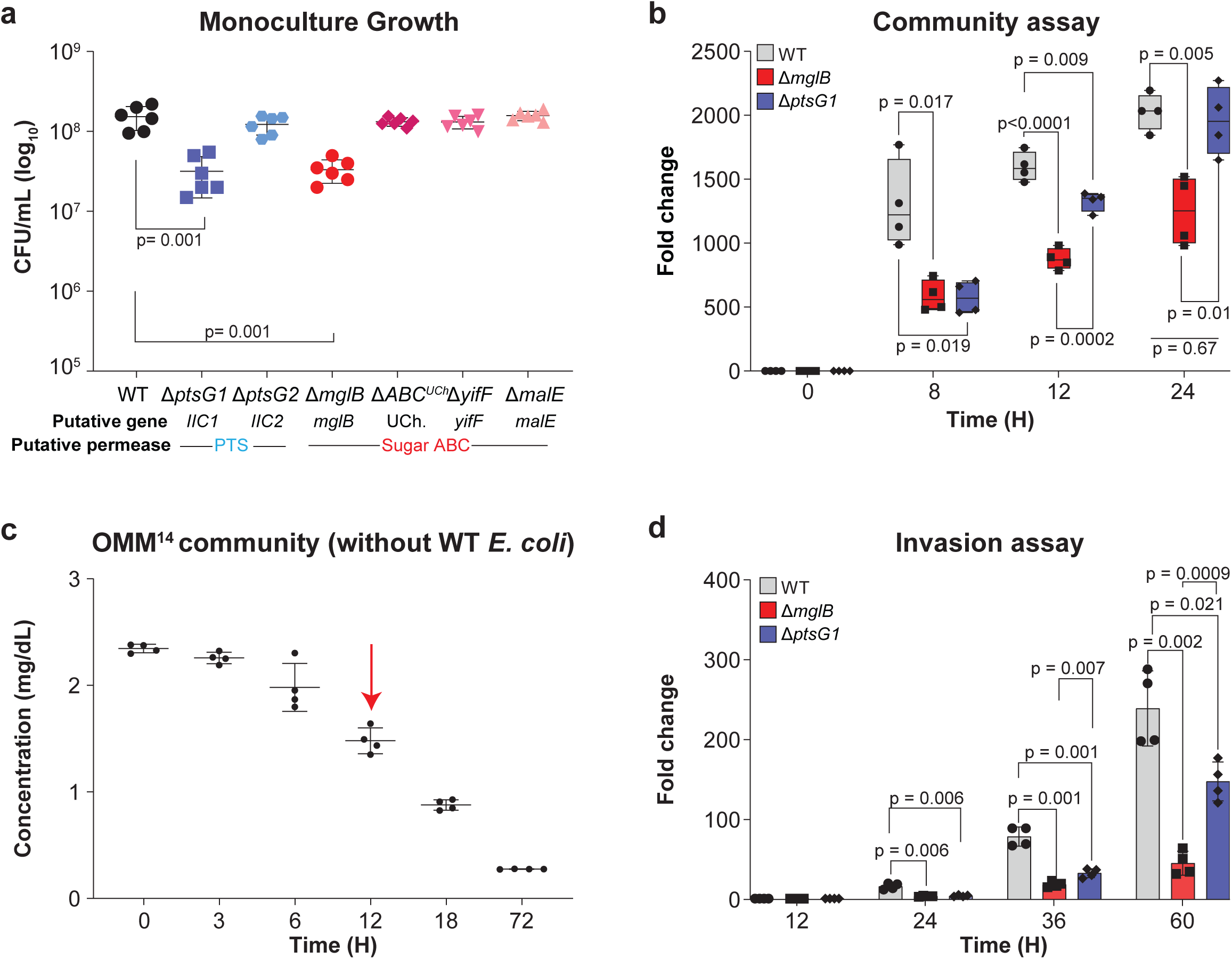
A sugar ABC transporter is necessary for ecological success and invasion of *E. coli* in a model community. **a)** Growth comparison measured as CFUs between wild-type (WT) *E. coli* and sugar transporter mutants cultured in BHI-NG medium supplemented with 0.25% GLU, measured at 24 h post-inoculation. Initial inoculum at 0 h was 10^5^ CFU/mL ± 0.3. **b)** Community assay showing fold change in density relative to 0 h for WT *E. coli* and its transporter mutants. WT *E. coli* and its transporter mutants were each grown separately as part of an OMM^15^ community containing 14 other members from the start (0 h) and monitored over 72 h in BHI-NG medium supplemented with 0.25% GLU. Fold changes were calculated from normalized qPCR Ct values (Ct/mL). The initial inoculum at 0 h was 10³ CFU/mL ± 0.3. **c)** Glucose concentration in culture supernatants of the OMM^14^ community over 72 h. Cultures were diluted 1:100 prior to measurement. Red arrow indicates the time point at which WT *E. coli* and its deletion mutants were introduced in the invasion assay shown in panel d. **d)** Invasion assay showing fold change in density of WT *E. coli* and its transporter mutants relative to the 12 h introduction time point. Each strain was introduced into a separate, pre-established OMM^14^ community at 12 h and monitored over the following 60 h in BHI-NG medium supplemented with 0.25% GLU. Fold changes were calculated from normalized qPCR Ct values (Ct/mL). The initial inoculum at 0 h was 10³ CFU/mL ± 0.3. For panel **a**, data are combined from two independent biological experiments with three technical replicates each (total n = 6). For panels **b–d** data are combined from two independent biological experiments with two technical replicates each (total n = 4). Statistical analyses for panels **a**, **b**, and **d** were performed using Brown–Forsythe and Welch’s ANOVA. *P* < 0.05 was considered statistically significant.

Our data revealed that transport systems for simple sugars are key determinants in microbial competition, and that they were more informative of growth in complex gut communities than central metabolism gene repertoires. Moreover, we find that a single genomic feature—possession of sugar ABC transporters—has outsized predictive power of complex gut microbiome dynamics, including the ability to invade a resident community.

## Discussion

Our study identifies sugar ABC transporters as genomic features with outsized predictive power over bacterial growth responses. This predictive capacity spans *in vitro* assays, *in vivo* supplementation with dietary sugars, and human microbiome data, establishing a single genomic trait as a key determinant of dietary sugar-mediated community dynamics. Because our findings were derived from diverse bacterial strains with variable central metabolism gene repertoires, we were able to rank sugar ABC transporters as the dominant fitness-driving feature in various models of gut microbial competition.

How is the competitive advantage conferred by sugar ABC transporters during sugar supplementation? One possibility is that increased sugar availability concentrates competition within the microbial community on sugars^4,16^, raising the ecological importance of high-affinity uptake systems like ABC transporters, even when absolute sugar concentrations are elevated. Usage of ABC and/or PTS transport systems varies between species and is regulated by substrate availability^19,21–23,43,44^. Sugar ABC transporters exhibit high substrate affinity and capacity for diverse sugar uptake, enabling efficient nutrient acquisition^45,46^ under a range of concentrations^19,20,47^. Additionally, different transporter systems may contribute at distinct stages of community dynamics in sugar-supplemented environments^47,48^. Our knockout experiments show that while both PTS and ABC transporters support early bacterial growth, only the ABC transporter is essential for sustained expansion and successful invasion within the community context (Fig. **5b** and **5d**). This suggests that although multiple nutrient uptake systems influence microbial fitness^19,21,47,48^, sugar ABC transporters provide a unique advantage when sugar competition is a dominant force shaping community structure.

Moreover, in HCT patients, whose microbiomes experience extreme antibiotic- and treatment-related microbiome disruption, which may sensitize them to effects of dietary sugars^3,40,49^, we found that genera with higher sugar ABC transporter counts had positive association with dietary sugar intake, consistent with sugar-induced expansion of such taxa. Interestingly, while not all genera with low transporter gene content declined, we observed a particularly strong negative association of *Prevotella* with dietary sugar intake, a genus typically lost in industrialized populations as a result of low-fiber diets and whose members lack sugar ABC transporters^2,14,31,32,50^. These findings demonstrate that sugar ABC transporter content is a genomic correlate of microbiome dynamics in response dietary sugars that extends to humans.

Taken together, the key outcome of this study is that sugar ABC transporters—a single genomic trait—consistently predict bacterial growth in sugar defined environments, across monoculture conditions, community invasion, *in vivo* expansion, and human microbiome responses to dietary sugars. The outsized predictive power of a single genomic feature described here is particularly notable given the complexity of microbial ecosystems, which are influenced by factors such as osmolarity, pH levels^29,30^, secondary metabolite production^33,51^ and cross-feeding^52^. As nutrient blocking is the prevailing ecological explanation for colonization resistance against pathogens^34,42^, we speculate that in diets defined by simple sugar intake, sugar ABC transporters are crucial mediators of this fundamental microbiome function. As such, our work also supports prioritization of functional traits over taxonomy in explaining microbiome dynamics^53–55^. Moreover, our results contribute to explaining the increase of Pseudomonadota, Bacillota, and pathobionts observed in dysbiosis and as a consequence of modern dietary shifts^3,11–14,31,32,50,56^, which often involve elevated consumption of refined sugars alongside reduced fiber intake. As global sugar intake increases, these findings demonstrate that specific gene content can be used to anticipate microbiome shifts, offering a predictive link between microbial gene content and microbiome responses to diet-driven ecological restructuring.

## Methods

### Strains and media

All bacterial strains used in this study are listed in Supplementary Tables 1 and 4. Starter cultures of human isolates were grown in Brain Heart Infusion broth without glucose (BHI-NG) (Bioworld), supplemented with 0.25% glucose (w/v) (Sigma-Aldrich). Starter cultures of the OMM^15^ community were also cultured in BHI medium under the same conditions, except A. *muris* and *S. danieliae*, which were cultured in Columbia agar medium (Remel) supplemented with 5% defibrinated sheep blood (Fisher Scientific). *A. muciniphila* and *B*. *caecimuris* were cultured in BHI medium supplemented with 0.24% GLU and 0.2% mucin (w/v) (Sigma-Aldrich). All bacteria were cultivated anaerobically at 37°C inside a Coy Model 2000 incubator in a type B anaerobic chamber (N2/CO2/H2, 95%/2.5%/5%) (Coy Laboratory Products, Inc., Grass Lake, MI, USA). Selective media for cloning purposes were prepared using LB broth or LB-agar, supplemented with antibiotics at the following final concentrations: 25 μg/mL chloramphenicol (Sigma-Aldrich) and 100 μg/mL ampicillin (Sigma-Aldrich).

### Cross-Phylum Genome Analysis of Sugar Transporters, Glycolytic Enzymes, and Pathways

Gut microbiome species were obtained from the BacDive database. Species were ranked by the number of completed genomes available in the NCBI database as of May 12, 2025. For each phylum, the top 20 species were selected for analysis, with *A. muciniphila* (the sole representative of Verrucomicrobiota) included. For each species, all available complete genomes were downloaded; if more than 20 genomes were available for a species, a random subset of 20 genomes was selected. To determine the counts of sugar transporters and glycolysis enzymes in bacterial genomes, genome annotation files were downloaded from NCBI. The genomes of *R. intestinalis*, *K michiganensis* AD9, and *E. coli* AD24 were annotated using PGAP^57,58^ provided by NCBI and these annotation files were used for genome searches. For the screening of sugar transporters, genome annotations were searched using the keywords “sugar” and “transporter.” Identified sugar transporters were then categorized into subgroups, including sugar ABC transporters, PTS transporters, and MFS transporters. Key glycolysis enzymes were selected from three primary glycolytic pathways: Embden-Meyerhof-Parnas (EMP), Pentose Phosphate Pathway (PPP), and Entner-Doudoroff (ED). For the screening of glycolytic enzymes, genome annotations were searched using the specified gene names (Supplementary Table 2). For hexokinase, alternative search terms such as “sugar kinase,” “glucokinase,” and “ROK family kinase” were employed. These selections provided comprehensive coverage of the different glycolytic pathways. For KEGG pathway analysis, KOALA^59^ was used to assign KEGG Orthology (KO) terms to proteins in deduplicated bacterial protein files. The annotated KOs were then used to reconstruct KEGG pathways in each genome via KEGG Mapper. Only complete glycolysis pathways were retained and considered functional for this study.

### CFU analysis

For CFU experiments, 5 mL of BHI-NG medium supplemented with 0.25% and 1% glucose (w/v) was inoculated with each bacterium to a final concentration of 10⁵ ± 0.3 CFU/mL, and cells were grown at 37 °C under anaerobic conditions without agitation. At various time points, cells were transferred to a 96-well flat bottom plate (Corning Costar), and serial dilutions ranging from 10^-1^ to 10^-7^ were performed in deoxygenated PBS. From these dilutions, 20 µL of sample was transferred to a BHI agar plate. Plates were incubated at 37 °C under anaerobic conditions, and CFUs were counted. Statistical analysis of gene presence/absence as predictor of CFUs was conducted with a linear mixed effects model using varying intercepts for biological repeats; the model was implemented in Python using the *statsmodels* library, with the following model equation for the regression of gene presence as predictors of CFUs at a given sugar concentration and time point: (CFU | time point, sugar concentration) ∼ ABC + PTS + MFS + RPI + PGM + KDGP + (1 | uniqex), where ‘uniqex’ represents the index of the biological repeat. Transporter gene presence was represented using binary indicators for ABC, PTS, and MFS. Glycolytic pathway genes were selected based on prevalence, retaining only those present in more than 25% but fewer than 75% of species. Binary indicators RPI, PGM, and KDGP correspond to ribose-5-phosphate isomerase, phosphoglucomutase, and 2-keto-3-deoxy-6-phosphogluconate aldolase, respectively.

### pH Experiments

For pH adjustment experiments (Supplementary Fig. 4), pH was measured using a glass double-junction electrode (Orion™ Star A111, Thermo Fisher Scientific), and media pH was titrated using 1 M NaOH to match across sugar concentrations.

### Growth measurements

For growth assays, BHI-NG medium was supplemented with 0.25% or 1% (w/v) glucose, fructose, sucrose, or galactose and inoculated with each bacterium to a final concentration of 10⁵ ± 0.3 CFU/mL in 250 µL of medium. Cultures were grown in a 96-well flat-bottom plate (Corning Costar) at 37°C under anaerobic conditions without agitation, using a Tecan Infinite Mplex plate reader. Prior to each optical density measurement, a 10-second double-orbital shaking step was performed. Optical density at 600 nm (OD₆₀₀) was recorded every 30 minutes for 72 hours.

### Bayesian models of sugar-genus associations and genome sugar transporter counts

The sequencing protocol for the16S rRNA gene sequencing for this dataset has been reported previosuly^3^. Assigned taxonomy were aggregated at the genus level, discarding amplicon sequence variants without genus identification. A relative abundance and prevalence threshold was then set to keep genera for analysis that were present in at least 10% of samples at relative abundance 0.01%, or that were present in any sample at 10% relative abundance or more. Of these 117 genera present after filtering, 91 had complete genomes on NCBI. To count sugar transporters of each type, we downloaded up to 100 randomly sampled genomes from each genus and counted the occurrence of “ABC”, “MFS”, and “PTS” in the feature products where “sugar” and “transporter” were already present. We reported means of these counts to approximate the count of sugar transporters of each type for each genus. “All sugar transporters” is a sum of these means.

Sugar-genus models. The associations of the 91 genera with sugar intake in the prior 2 days were quantified with individual Bayesian univariate models with the following definition,

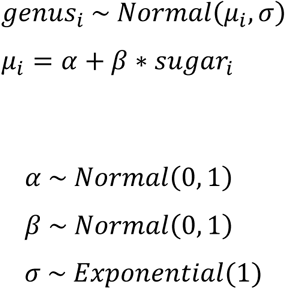

for sample *i* =1009 for each of the 91 genera. The relative abundance matrix of samples and genera used in the analysis was centered-log ratio (CLR)-transformed^60^ the predictors and outcome were standardized, and a Hamiltonian Monte Carlo sampler was used to extract 4000 samples (across 4 chains) from the posterior distributions (R v4.3.2, rstan v2.32.6, Stan v2.32.2). Posterior distributions for the *β* parameter associated with the sugar predictor were summarized by plotting the mean value and the 66% and 95% credible intervals.

The relationship between sugar transporter counts and each genus’s association with sugar was investigated in two ways. Firstly, for each sugar transporter type (and all sugar transporters), a Spearman rank correlation was performed between the mean count of each sugar transporter type and the mean values from the sugar parameter of each Bayesian sugar-genus model. Secondly, a Bayesian linear regression was visualized following the model definition,

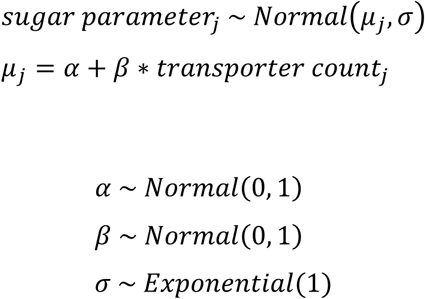

for *j* = 91 genera. The sugar parameters existed on the z-score scale already, and sugar transporter count was normalized by its maximum before sampling. For each genus, *µ_j_* was simulated over the range of sugar transporter values using 4000 posterior samples (1000 x 4 chains), and the mean of *µ_j_* was plotted with a 95% credible interval.

### HCT patient dataset

The previously published dataset^3^ comprised 158 patients undergoing HCT, 1009 longitudinally collected fecal samples on which microbiome profiling was conducted with 16S rRNA gene sequencing, and 9419 meals with dry weight in grams of each food item consumed (as indicated in quarter-portions by the patient and verified by a nurse). We used the per-food-item data to sum up the diet (grams of each food item) of each patient every day, and matched this dietary information to microbiome samples by taking the mean of the day sums from the 2 days prior to each microbiome sample (e.g., a sample collected on day 3 is associated with the mean of the diet sums of day 1 and day 2). Macronutrient composition of each food item was collected from the USDA FNDDS database, where each of our food items represented the highest-resolution 8-digit food code. We could then calculate mean sugar intake in grams over the prior 2 days for every microbiome sample.

### *In vivo* experiments

#### Mice

Mice were kept on a 12h light/dark cycle with ad libido access to autoclaved rodent chow and drinking water. Offspring of germ-free C57BL/6Tac mice housed in sterile isolators were used for all experiments.

Animal care and experimentation were consistent with NIH guidelines and approved by the Institutional Animal Care and Use Committee (IACUC) at New York University School of Medicine

#### OMM^15^ colonization

*T. muris* and *M. intestinale*, strains were purchased from DSMZ, while the remaining strains were generously provided by Xiaomin Xiao and Ken Cadwell (NYU School of Medicine/University of Pennsylvania). Strains were cultured under anaerobic conditions as described above, combined at equal CFU ratios 10^8^ ± 0.3, and the resulting mixture was stored in 20% glycerol at -80°C for further use.

Eight-week-old male and female germ-free C57BL/6Tac mice, housed in sterile isolators, were used for colonization. On the day of colonization, the bacterial mixture was thawed and kept on ice until inoculation. Mice were inoculated via oral gavage and enema using a 1 mL syringe fitted with an 20ga, 38mm flexible plastic rodent feeding tube (Instech, Cat# 50-475-764). A second dose of the bacterial mixture was administered on day 2 post-initial inoculation.

To allow for recovery and minimize stress, breeders were set up on day 7 following the secondary inoculation. Vertically colonized mice were used for all experiments.

#### *In vivo* high sugar challenges

Mice were randomly assigned to treatment groups and provided with either 10% glucose (Sigma-Aldrich), 10% fructose (Sigma-Aldrich), or 10% sucrose (Sigma-Aldrich) dissolved in filtered drinking water. Control mice received filtered drinking water without supplementation. All sugar solutions and control water were sterilized using a SteriCup 0.22 µm filter (Millipore) before administration.

Body weight and fluid intake were monitored throughout the experiment, and fluids were replaced every three days to maintain freshness and cleanliness. Fresh fecal samples were collected at baseline (day 0) and on day 14, immediately stored on dry ice, and subsequently kept at -80°C for long-term storage. Final body weight measurements were recorded on day of sacrifice (day 14).

#### Murine fecal DNA Extractions

Fecal weights were recorded prior to DNA extraction. DNA was extracted from fecal samples using the Quick-DNA Fecal/Soil Microbe MiniPrep Kit (Zymo Research, D6010) following the manufacturer’s protocol.

#### Mouse intestine dissections

Mice were sacrificed by cervical dislocation, and the entire intestinal tract, spanning from the gastroduodenal junction to the rectum, was carefully removed as an intact piece. The intestine was divided into segments as follows: Small intestine: Defined from the gastroduodenal junction to the ileocecal junction; Cecum: Isolated as a distinct segment; Colon: Defined as the section extending from 1 cm distal to the cecal-colonic junction to 1 cm proximal to the anus. The lengths of each segment were measured by aligning them along a ruler on a flat surface to ensure accuracy.

### DNA extraction and Quantitative PCR of bacterial 16S rRNA genes

DNA was extracted from *in vitro* community and invasion assay using the Fecal/ soil microbe MiniPrep Kit (Zymo Research, D6010) following the manufacturer’s protocol. Quantitative PCR was performed as described previously^61^, with 1 μL of the extracted DNA used for qPCR reactions. Strain-specific 16S rRNA primers (Supplementary Table 8) at 10 μM concentration were employed, with SYBR FAST Roche LightCycler 480 2X qPCR Master Mix. The qPCR cycling conditions included a pre-incubation step at 95°C for 3 minutes, followed by amplification with 3 s at 95°C, 10 s at 60°C, and 20 s at 72°C, and a melting curve of 5 s at 95°C and 1 min at 65°C for a total of 40 cycles. Fold changes in the density of each bacterial strain across time points or experimental conditions were calculated directly from normalized qPCR Ct values. Absolute 16S rRNA gene copy numbers were not determined, as our analysis focuses on relative changes within the same strain. Ct values were normalized to the volume of culture (*in vitro*) or fecal weight (*in vivo*) for fold change calculations. The final fold change was determined using the equation

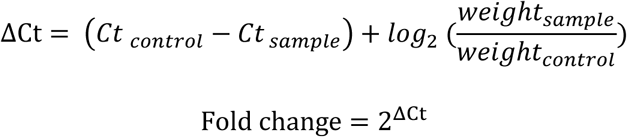

### *E. col*i MT1B1 mutants’ construction

All mutants generated in this study are reported in Supplementary table 5. Deletion mutants were generated as described previously in^62^. Briefly, plasmid pKD46 was introduced into wild-type *E. coli* MT1B1 via electroporation. A chloramphenicol resistance cassette (cat) was PCR-amplified from plasmid pKD3 using primers containing ∼100 bp homology arms flanking the target gene. The resulting PCR product was electroporated into *E. coli* MT1B1 harboring pKD46. Successful recombinants were selected on chloramphenicol-containing media, and gene deletions were confirmed by colony PCR. Plasmids and primers used are listed in Supplementary Tables 6 and 7.

### *In vitro* community experiments

OMM^15^ starter cultures were grown in their respective media as described previously. For community experiments, 500 mL of BHI-NG medium supplemented with 0.25% glucose (w/v) was inoculated with each bacterium to a final concentration of 10^3^ ± 0.3 CFU/mL. The cultures were grown at 37°C under anaerobic conditions without agitation. At defined time points, 20 mL aliquots were collected, centrifuged, and the resulting pellets were stored at −80 °C for subsequent processing and quantitative PCR analysis.

### Glucose measurements

To quantify free glucose concentrations in OMM^14^ (excluding *E. coli*) and OMM^15^ communities over time, cultures were set up as described above. At defined time points 1 ml samples were collected, centrifuged at 8,000 × *g* for 5 minutes, and the supernatants were filtered (0.22 µm). Glucose levels in the cleared supernatants were measured using a glucose colorimetric detection kit (Invitrogen, EIAGLUC), following the manufacturer’s protocol. Absorbance was read at 560 nm using a plate reader (Tecan Infinite Mplex), and glucose concentrations were calculated based on a standard curve generated in parallel.

### Invasion assay

The OMM^14^ community excluding *E. coli* was cultured as described above in BHI-NG medium supplemented with 0.25% GLU under anaerobic conditions at 37°C. After 12 hours, when glucose concentrations began to decline, wild-type *E. coli* MT1B1 or its respective deletion mutants were inoculated separately into established OMM^14^ cultures at a final concentration of approximately 10^3^ ± 0.3 CFU/mL. Cultures were maintained under the same anaerobic conditions, and 20 mL samples were collected at designated intervals, immediately stored at -80°C, and subsequently analyzed by quantitative PCR to determine bacterial population dynamics.

### Data analysis and figures

Data was analyzed using Graphpad Prism 9, python (v.3.8.9) and R (v.4.1.2). Figures were generated using Adobe Illustrator CC (Adobe Inc.). The statistical analysis varied for different datasets and details on the statistical methods are reported in the figure legends.

## Acknowledgements

This work was in part funded by an NIH/NIAID DP2 award (DP2AI164318) to J.S., NIH/NCI R01CA269617 to Catherine Diefenbach and J.S., and a Gabrielle’s Angel Foundation Award to J.S.. F.M is supported by a Helen Hay Whitney Fellowship. We furthermore thank the NYU Langone Health core facilities, including High Performance Computing, Gnotobiotics Core Facility, Genome Technology Center (RRID: SCR_017929), partially supported by the Cancer Center Support Grant P30CA016087 at the Laura and Isaac Perlmutter Cancer Center. We thank Xiaomin Xiao and Ken Cadwell for providing OMM^15^ strains and Baerbel Stecher for advice setting up the OMM^15^ colony.

## Author contributions

H.M. and J.S. conceived the study. H.M. performed the experiments with support from A.P.S,

F.M. performed the *in vivo* experiments, supported by E.K.N. C.D. performed the genomic comparison analyses. W.J. performed the human microbiome analyses. H.M., C.D., W.J., C.Z. and J.S. performed the computational analyses, and H.M., C.D., W.J., and J.S. interpreted the output and the data. H.M., C.D., W.J., A.P.S., and J.S. prepared the figures, and H.M. and J.S. wrote the manuscript, with contributions of methods sections by C.D., W.J. and F.M., and feedback from all other authors.

## Code Availability

Scripts for sugar transporter counting from NCBI, KEGG analysis, and Bayesian statistical models can be accessed in our GitHub repository (https://github.com/wjogia/microbiome-sugar-transporters).

---

**Extended Data Fig. 1.**
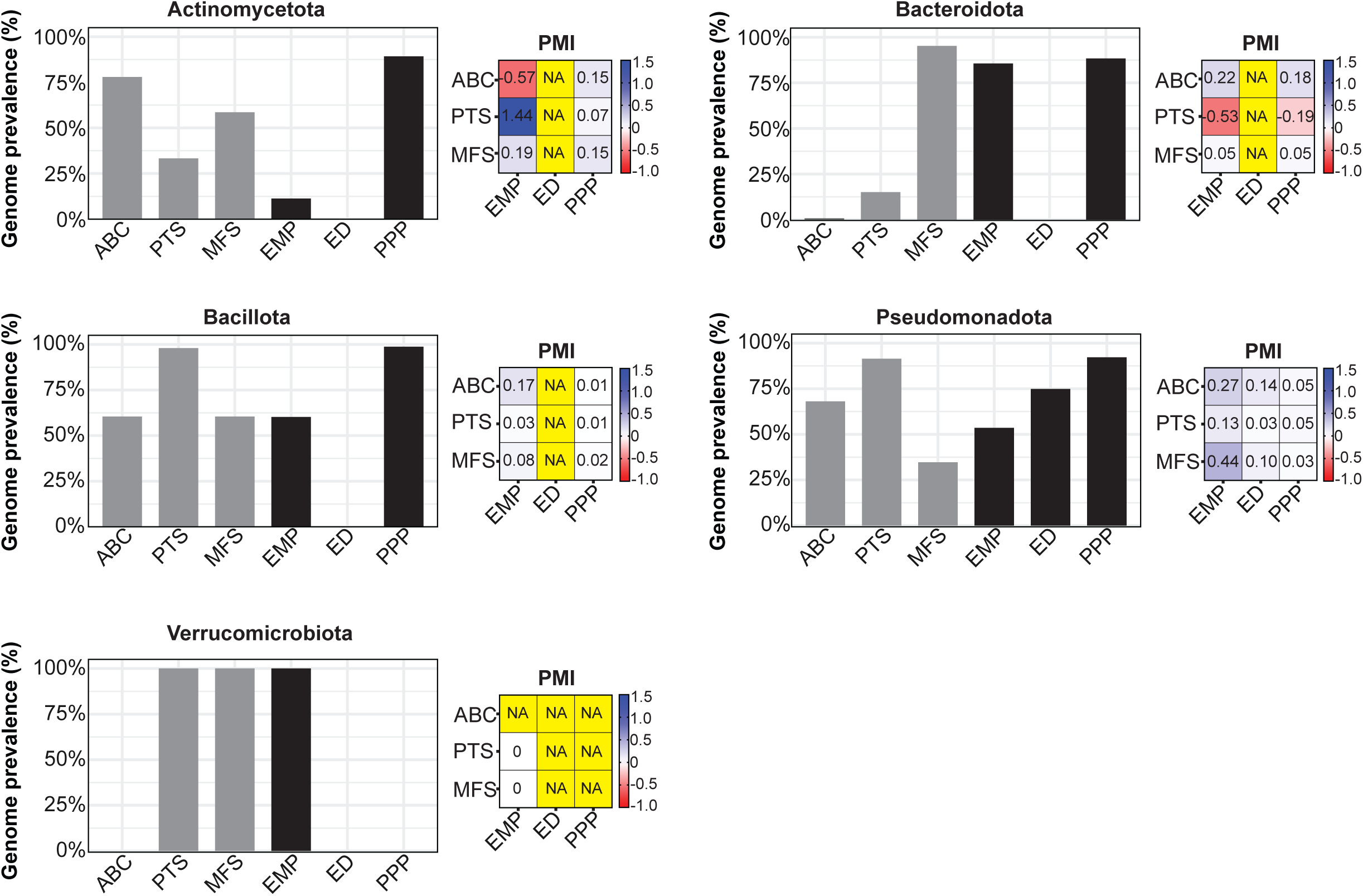
Sugar metabolism gene profiles reveal intra-phyla functional diversity in gut bacteria. Bar plots show the genome prevalence of sugar transporter types (ABC, PTS, MFS) and glycolytic pathways (EMP, ED, PPP) across bacterial phyla, highlighting functional differences within phyla. Heatmaps illustrate pairwise mutual information (PMI) between transporter and pathway features, capturing phylum-specific patterns. “NA” indicates missing PMI values due to the absence of one or both features.

**Extended Data Fig. 2.**
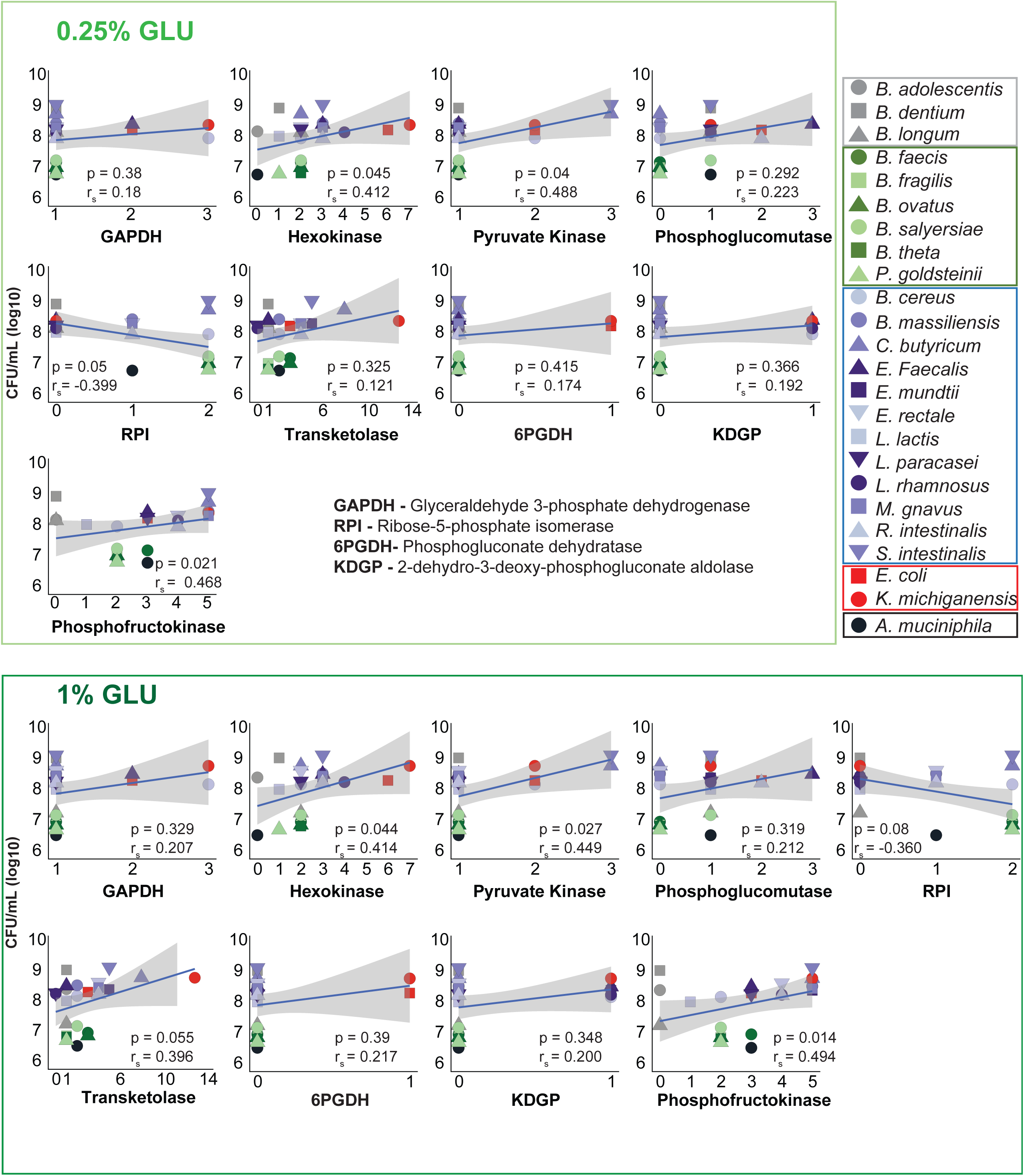
Growth correlates poorly with glycolytic gene profiles across EMP, PPP, and ED pathways. Spearman correlation (r_s_) analysis of the colony-forming units (CFUs) and genes involved in glycolytic pathways for 24 bacterial strains. Bacteria were cultured in BHI-NG medium supplemented with either 0.25% or 1% GLU, and samples were collected 24 hours post-inoculation. Each dot represents mean CFU of the indicated species from two independent experiments (n = 6). *P-*values are derived from Spearman correlation. Blue lines show linear regression fits; shaded areas indicate 95% confidence intervals.

**Extended Data Fig. 3.**
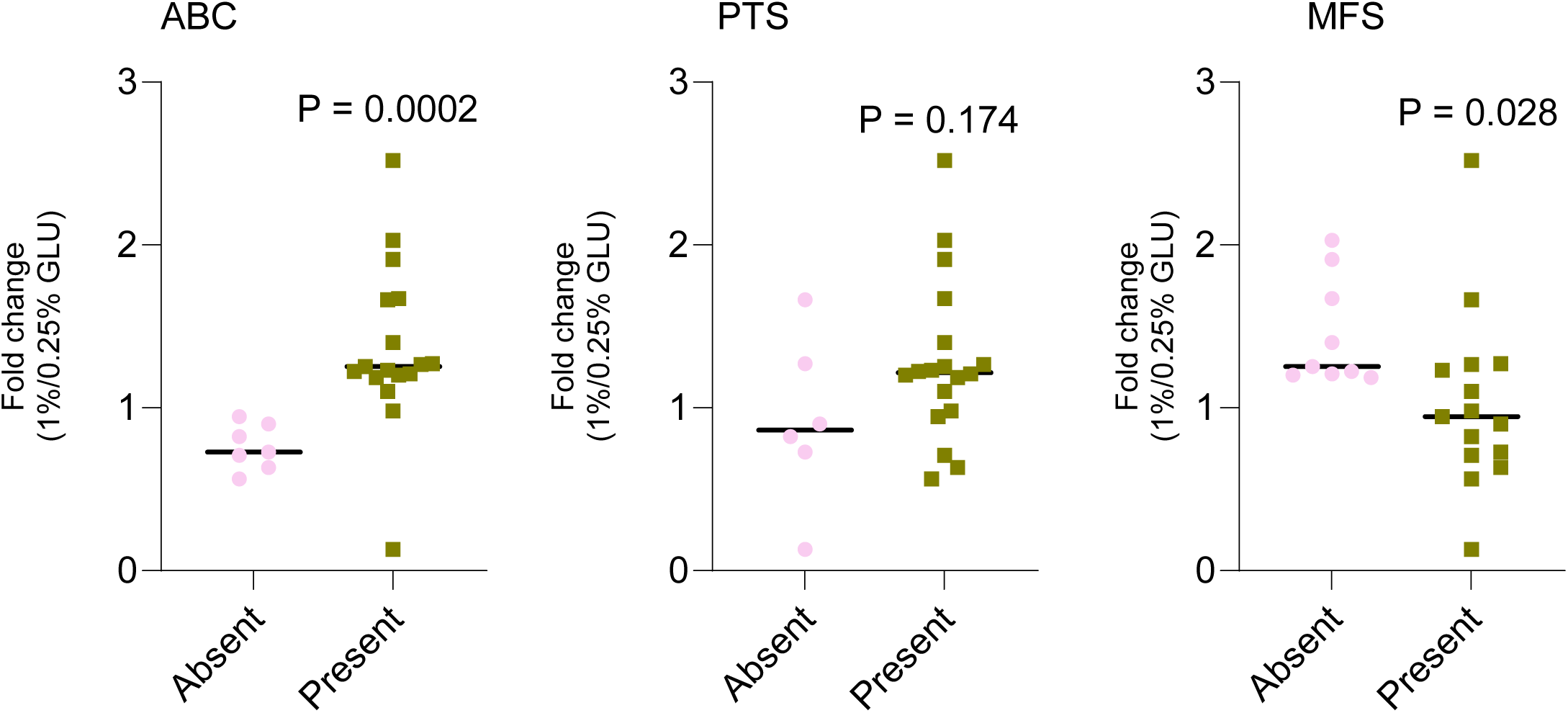
Sugar transporter presence distinguishes bacterial growth responses to glucose. Analysis of bacterial growth differences (fold change) based on the presence or absence of sugar transporters. Each data point represents the mean fold change in colony-forming units (CFUs) between 1% and 0.25% glucose (GLU) conditions at 24 hours for an individual strain, derived from two independent experiments (n = 6). Bacteria were cultured in BHI-NG medium supplemented with either 0.25% or 1% GLU, and samples were collected 24 hours post-inoculation. Statistical analysis was performed using an unpaired t-test with Welch’s correction, p < 0.05 was considered statistically significant.

**Extended Data Fig. 4.**
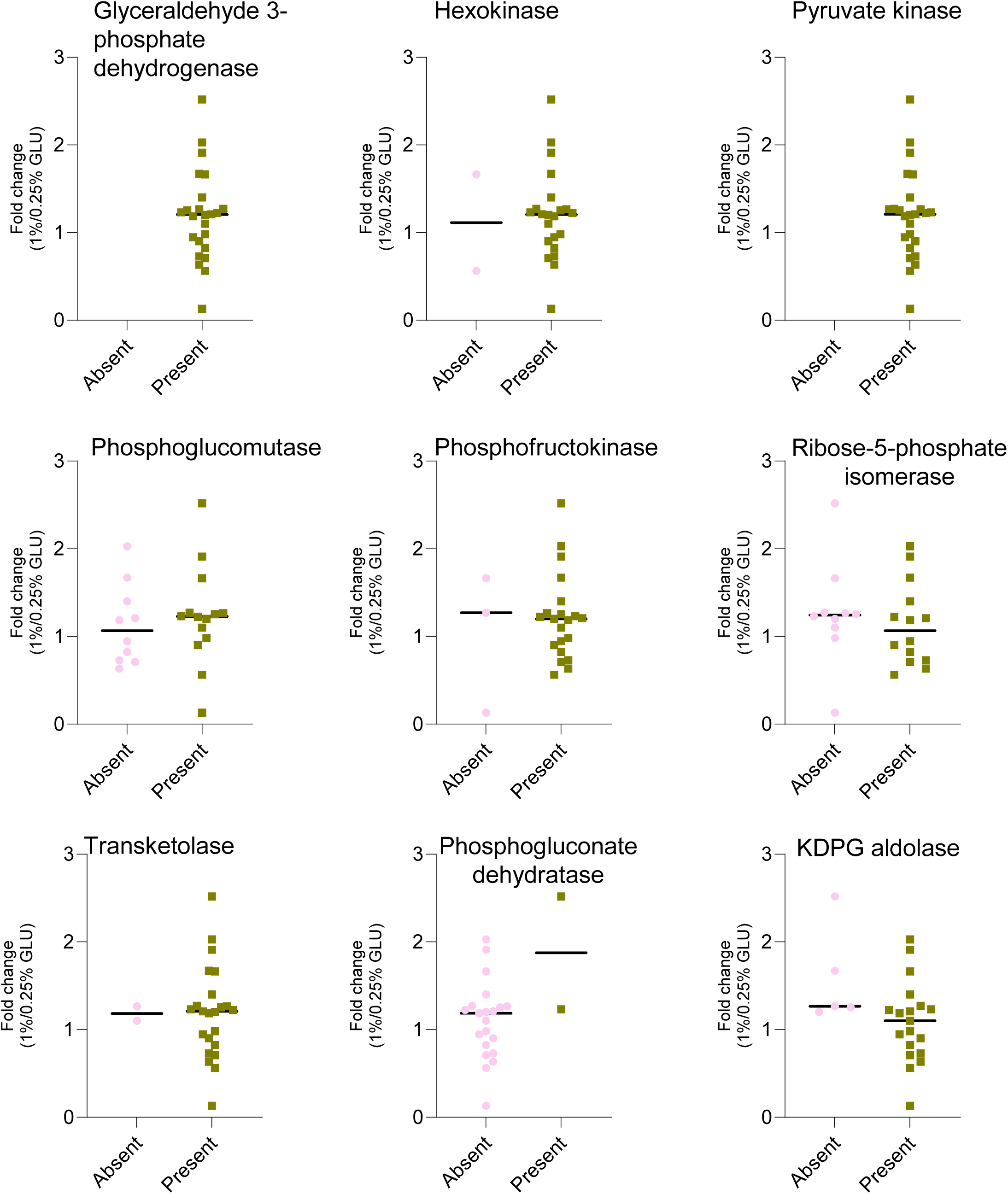
Bacterial growth responses to glucose are not determined by glycolytic enzyme presence. Analysis of bacterial growth differences (fold change) based on the presence or absence of enzymes involved in glycolytic pathways. Each data point represents the mean fold change in colony-forming units (CFUs) between 1% and 0.25% glucose (GLU) conditions at 24 hours for an individual strain, derived from two independent experiments (n = 6). Bacteria were cultured in BHI-NG medium supplemented with either 0.25% or 1% GLU, and samples were collected 24 hours post-inoculation. Statistical analysis was performed using an unpaired t-test with Welch’s correction, p < 0.05 was considered statistically significant. No significant differences were observed.

**Extended Data Fig. 5.**
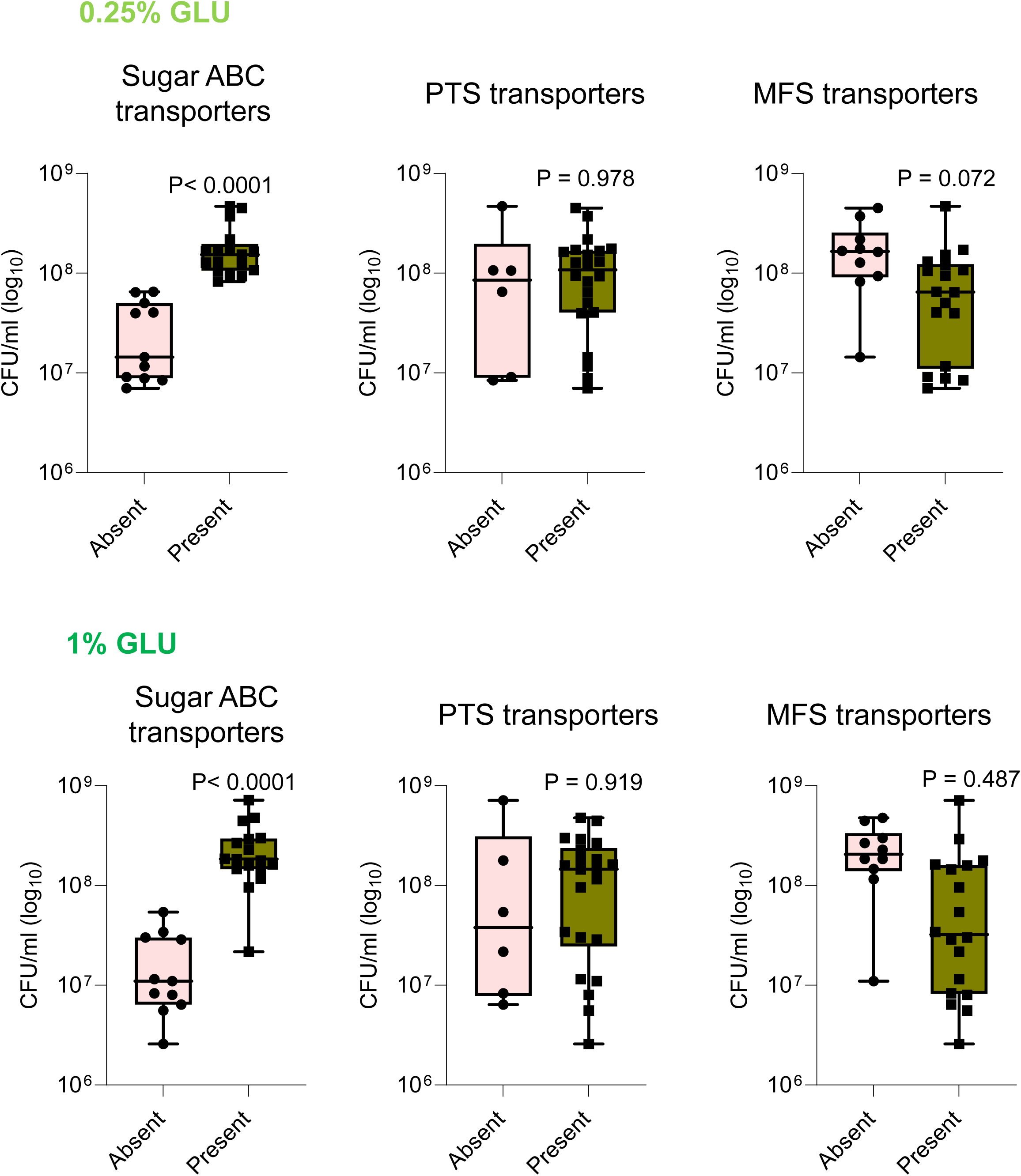
Strains encoding sugar ABC transporters show superior growth under varying glucose levels. Bar plots illustrating differences in growth among bacteria from Figures 1f and 2e, when categorized by the presence or absence of the indicated transporters. Each dot represents the average CFU of an individual species from two independent experiments (n = 6). Bacteria were cultured in BHI-NG medium supplemented with 0.25% or 1% glucose (GLU) and harvested at 24 hours post inoculation. Statistical analysis was performed using unpaired *t*-test with Welch’s correction. *p* < 0.05 was considered statistically significant.

**Extended Data Fig. 6.**
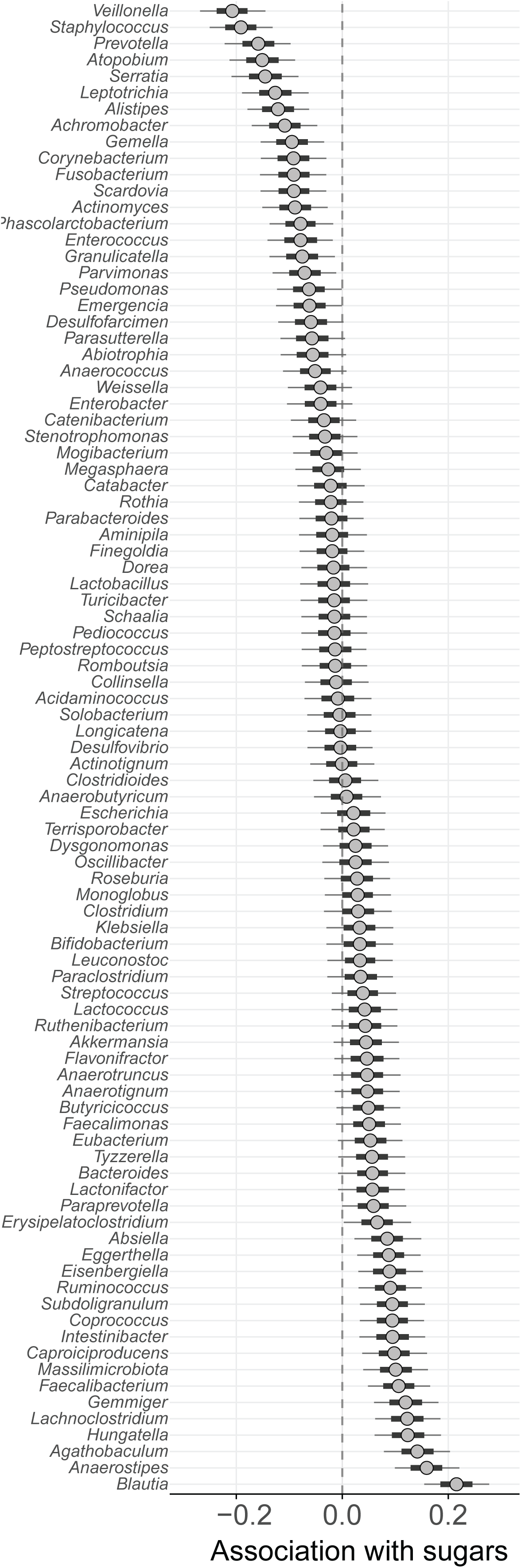
Bayesian inference of sugar intake effects on gut microbial genera in HCT patients. Genus–sugar associations are shown for all genera included in the HCT patient microbiome analysis. Bayesian linear regressions were performed between prior 2-day sugar intake and the relative abundance of each genus (n = 1009 samples, N = 158 patients, including all 91 genera). Posterior means for the effect of sugar are plotted with 66% and 95% credible intervals.

**Extended Data Fig. 7.**
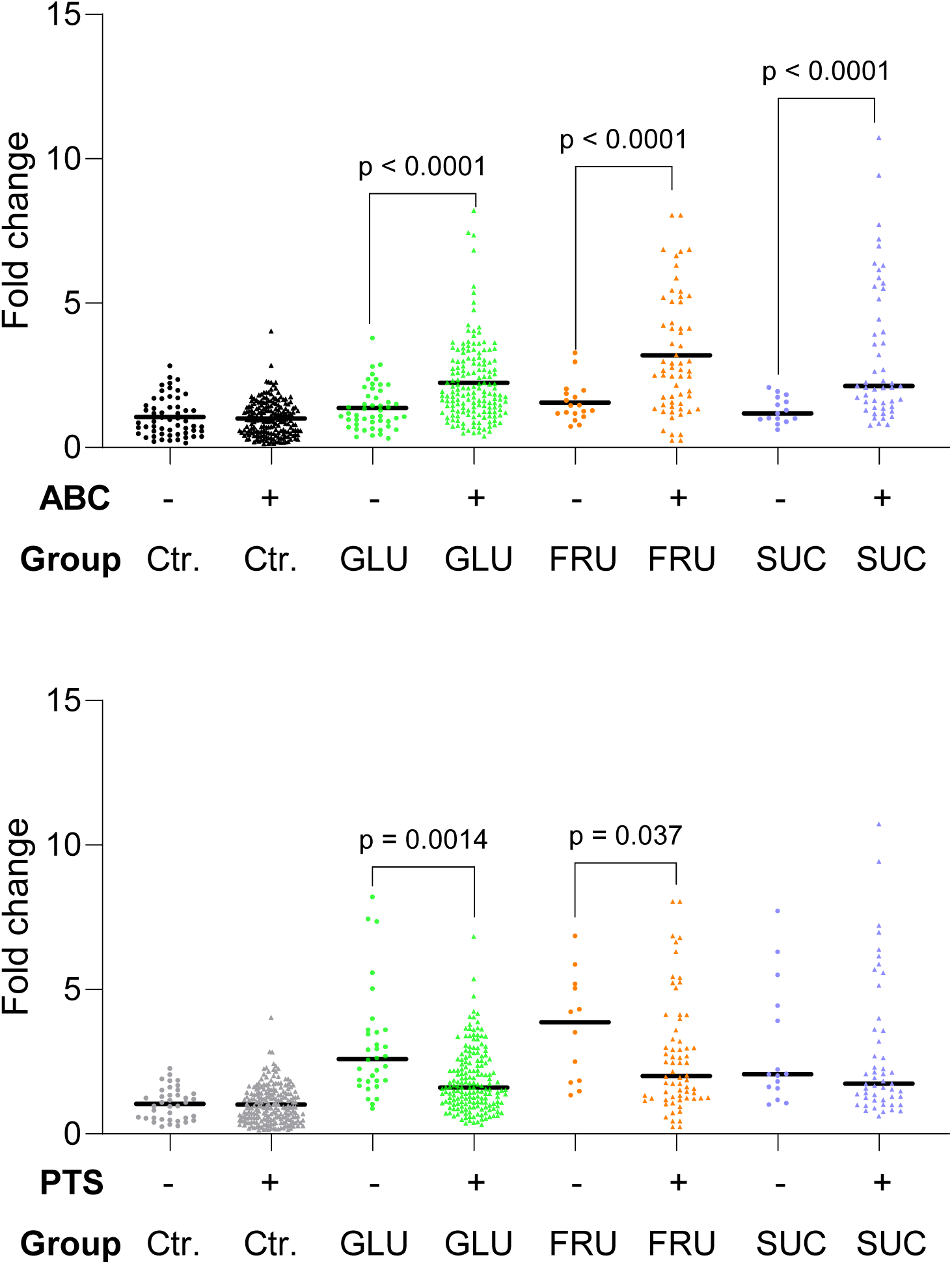
Transporter-specific growth responses of gut bacteria to dietary glucose, fructose, and sucrose. Fold change from fecal samples of C57BL/6 mice colonized with OMM^15^ species, with *ad libido* access to regular drinking water (control, Ctr.), or drinking water supplemented with 10% glucose (GLU), 10% fructose (FRU), and 10% sucrose (SUC) water, categorized by the presence (+) or absence (–) of ABC or PTS transporters. Each dot represents the mean fold change (from day 0 to day 14) for an individual bacterial species. Data were combined from 3 independent experiments (n = 10 control female mice, n = 9 control male mice, n = 7 10% GLU female mice, n = 9 10% GLU male mice, n = 3 10% FRU female mice, n = 3 10% FRU male mice, n = 3 10% SUC female mice, and n = 2 10% SUC male mice). Statistical analysis was performed using an unpaired t-test with Welch’s correction. P < 0.05 is considered statistically significant.

## Supplementary Information

**Supplementary Figure. 1.**
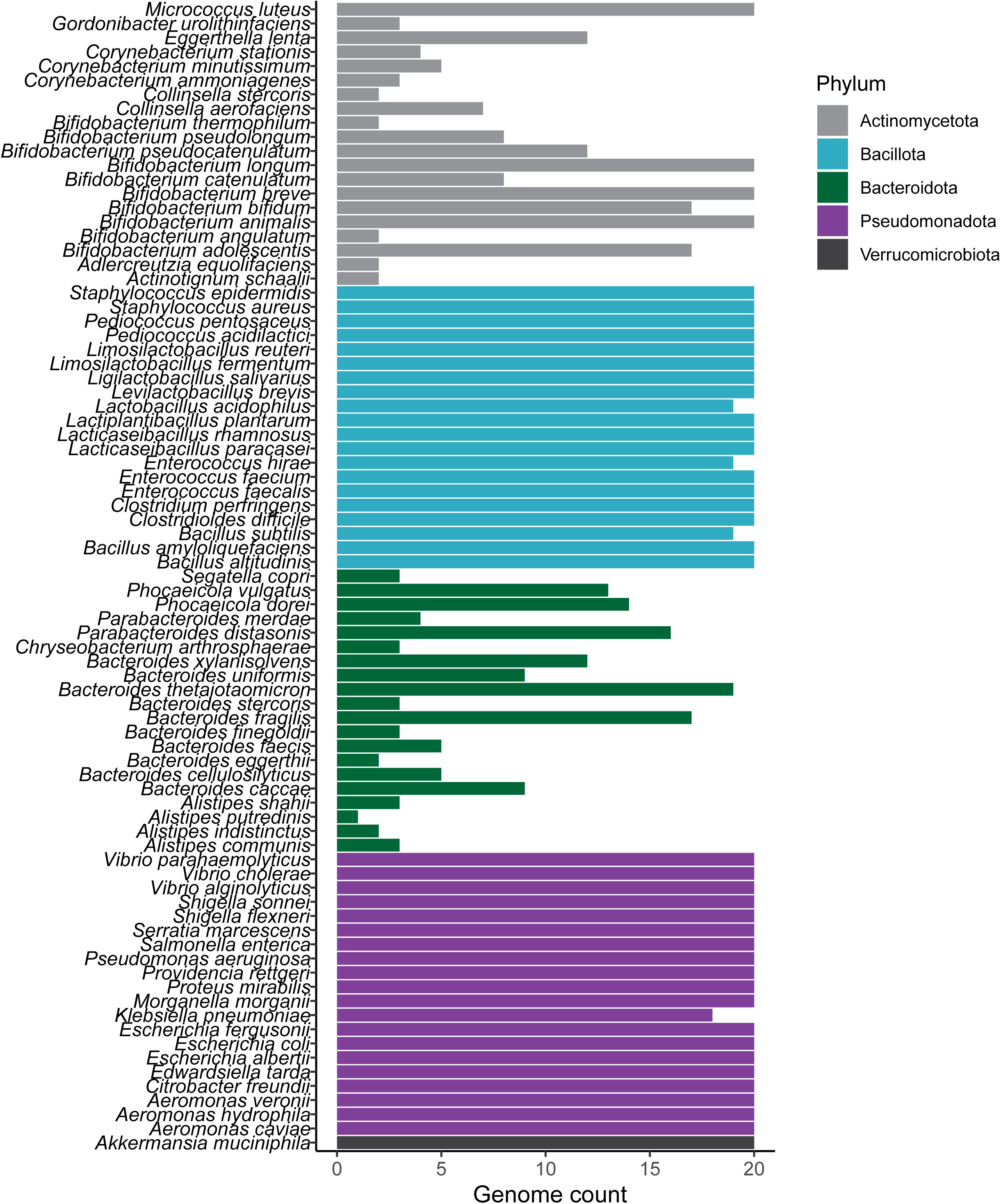
List of genomes used in Fig. 1a. For each phylum (excluding Verrucomicrobiota), 20 species were selected. For species with more than 20 fully sequenced genomes available, a maximum of 20 genomes were randomly chosen. For species with fewer than 20 sequenced genomes available, all available genomes were included.

**Supplementary Figure 2.**
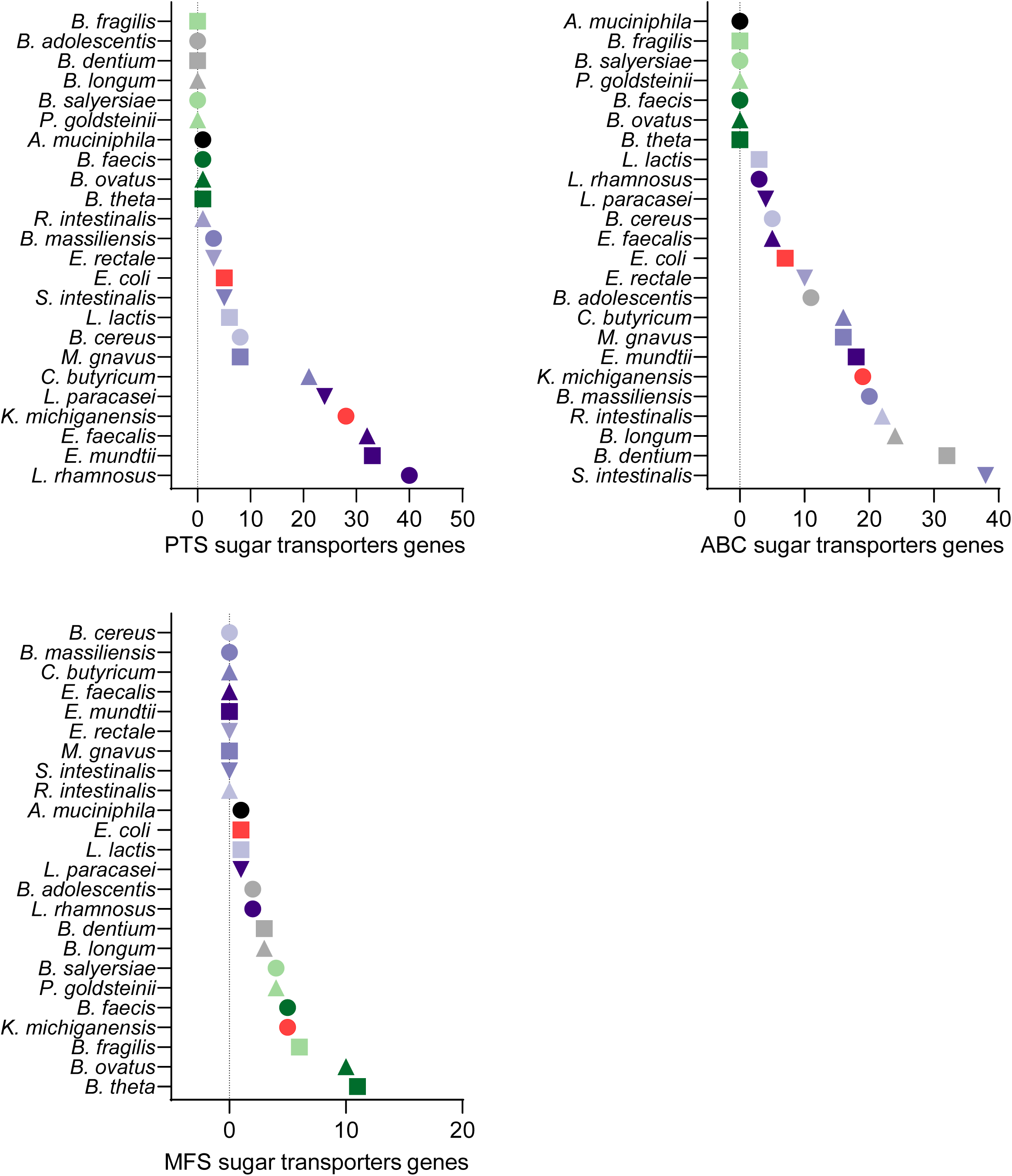
Enumeration of genes involved in ABC, PTS and MFS sugar transporters of the indicated bacteria.

**Supplementary Figure 3.**
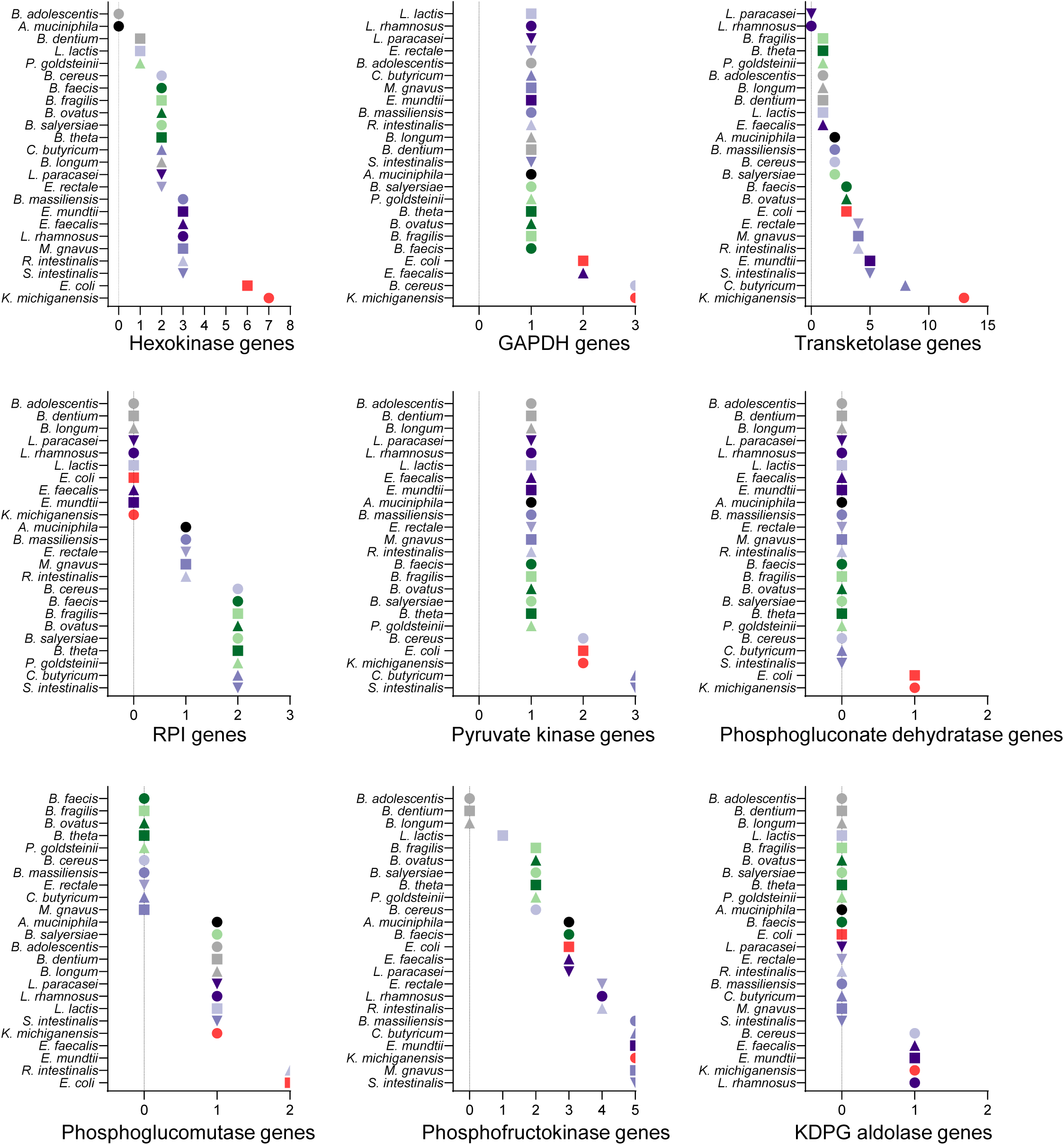
Enumeration of genes involved in the glycolytic pathways of the indicated bacteria.

**Supplementary Figure 4.**
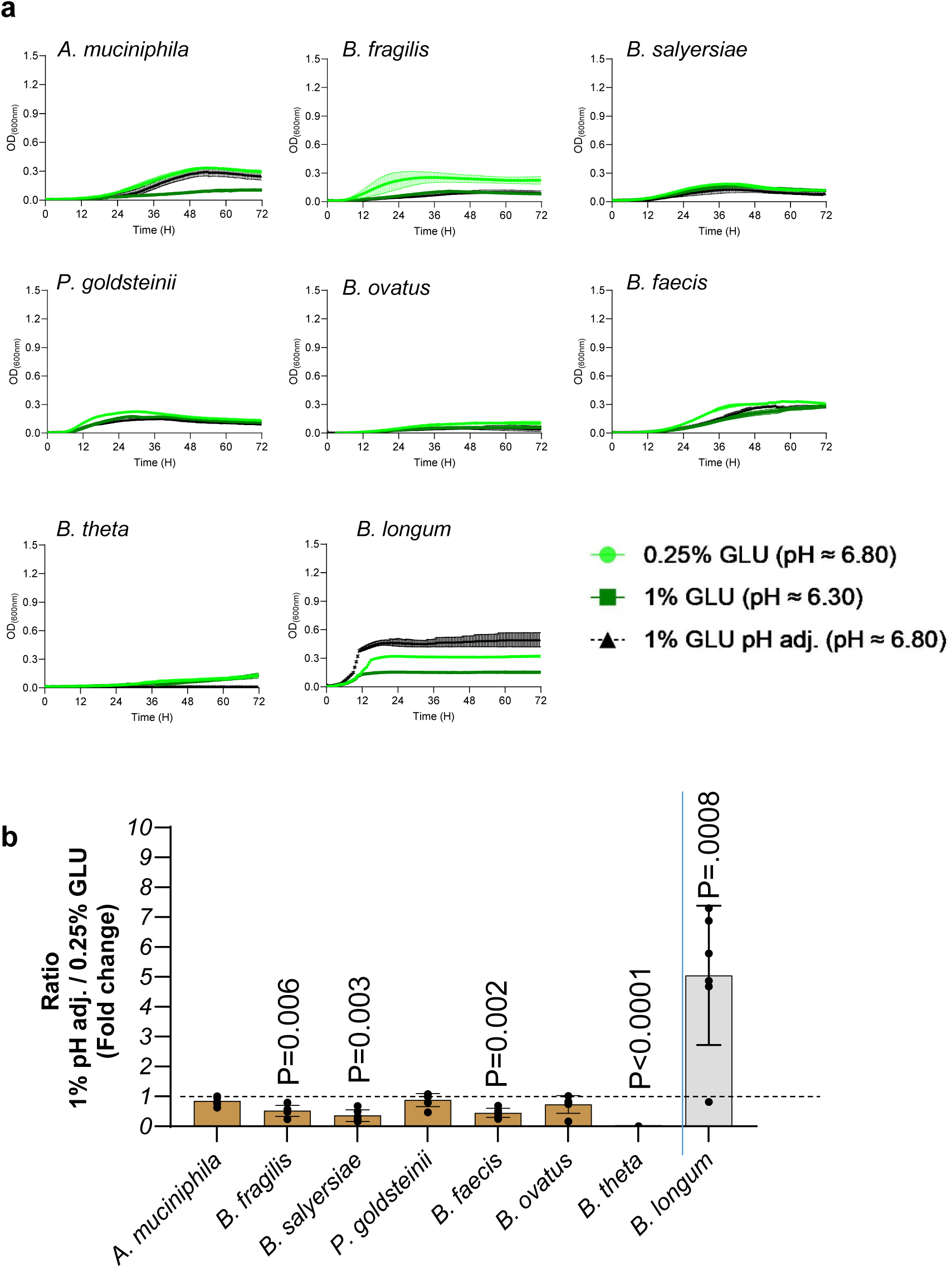
**a)** Analysis of the growth (OD_600_) of the indicated strains. Growth was monitored in BHI-NG medium supplemented with 0.25%, 1% and 1% (pH adjusted to 0.25% medium) glucose. Graphs represent mean ± SD from two independent experiments (n = 6). **b)** The fold change between the CFUs obtained at 1% pH adjusted and 0.25% glucose at 48 hours for each indicated species, derived from two independent experiments (*n* = 6). Bacteria were cultured in BHI medium supplemented with 0.25% or 1% pH adjusted glucose and harvested at 24 hours post inoculation. Data are presented as mean ± SD (with individual data points). Statistical analysis was performed using a two-tailed one-sample ratio t-test against a fold change of 1. *P* < 0.05 was considered statistically significant. The dashed line represents a fold change of 1, indicating no difference in growth between the two glucose concentrations.

**Supplementary Figure 5.**
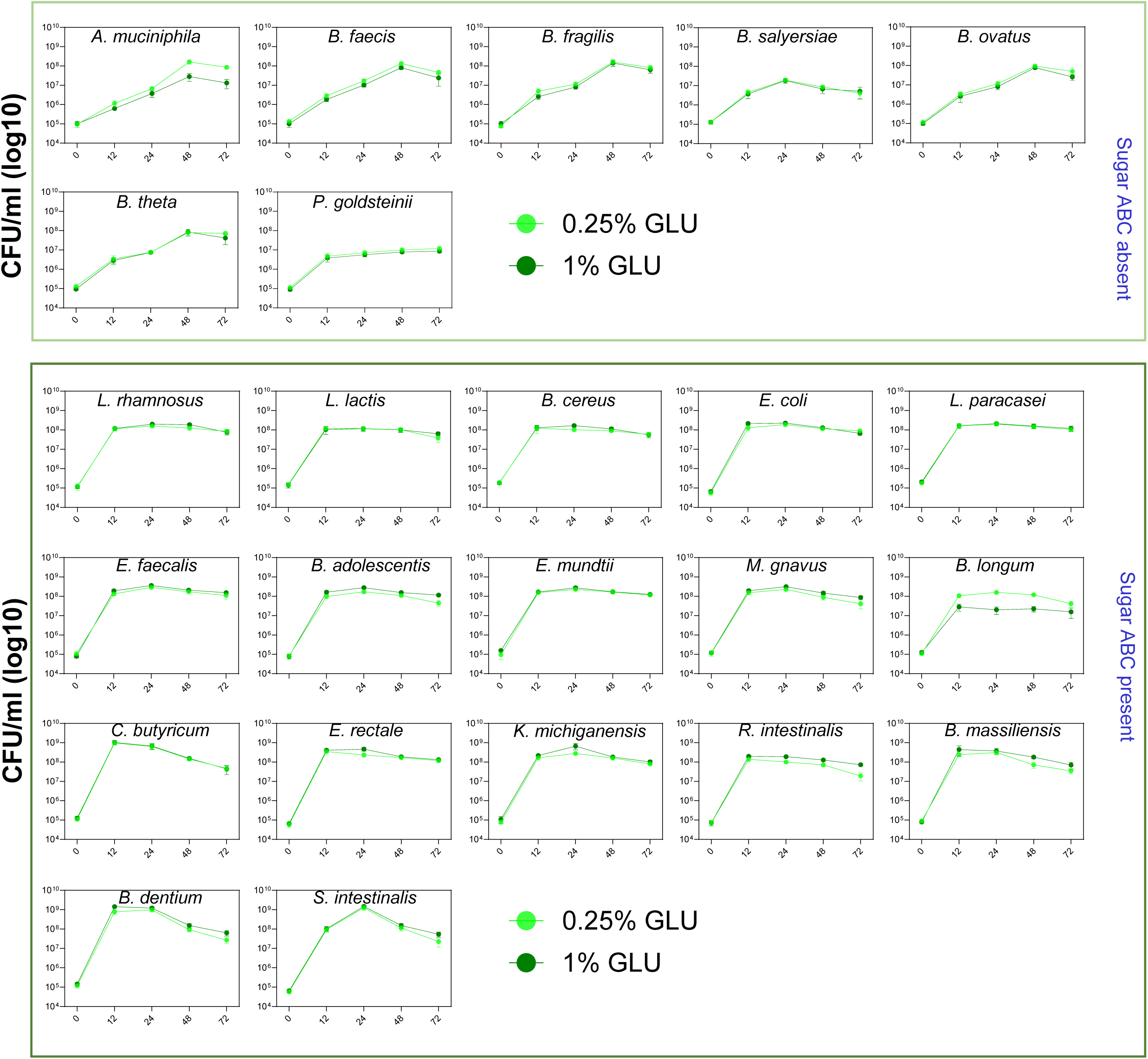
Growth analysis of the indicated bacteria. Bacteria were cultured in BHI-NG medium supplemented with 0.25% or 1% glucose (GLU) and harvested at 12, 24, 48 and 72 hours post inoculation and colony-forming units (CFU) were counted. Initial inoculum at 0 hour was 10^5^ CFU/mL ± 0.3 Graphs represent mean ± SD from two independent experiments (n = 6).

**Supplementary Figure 6.**
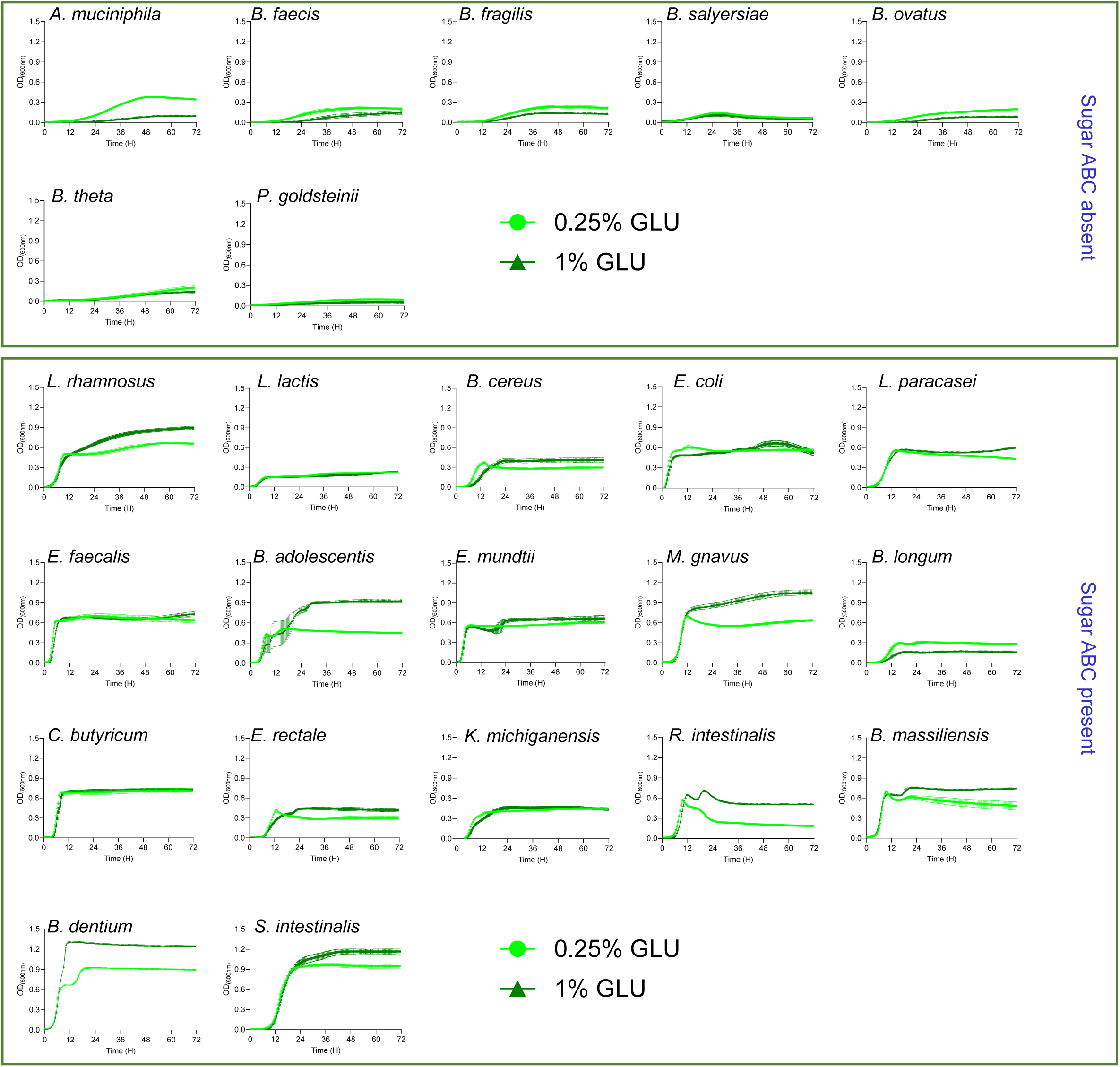
Analysis of the growth (OD_600_) of the indicated strains. Bacteria were cultured in BHI medium-NG supplemented with 0.25% and 1% glucose (GLU). Graphs represent mean ± SD from two independent experiments (n = 6).

**Supplementary Figure 7.**
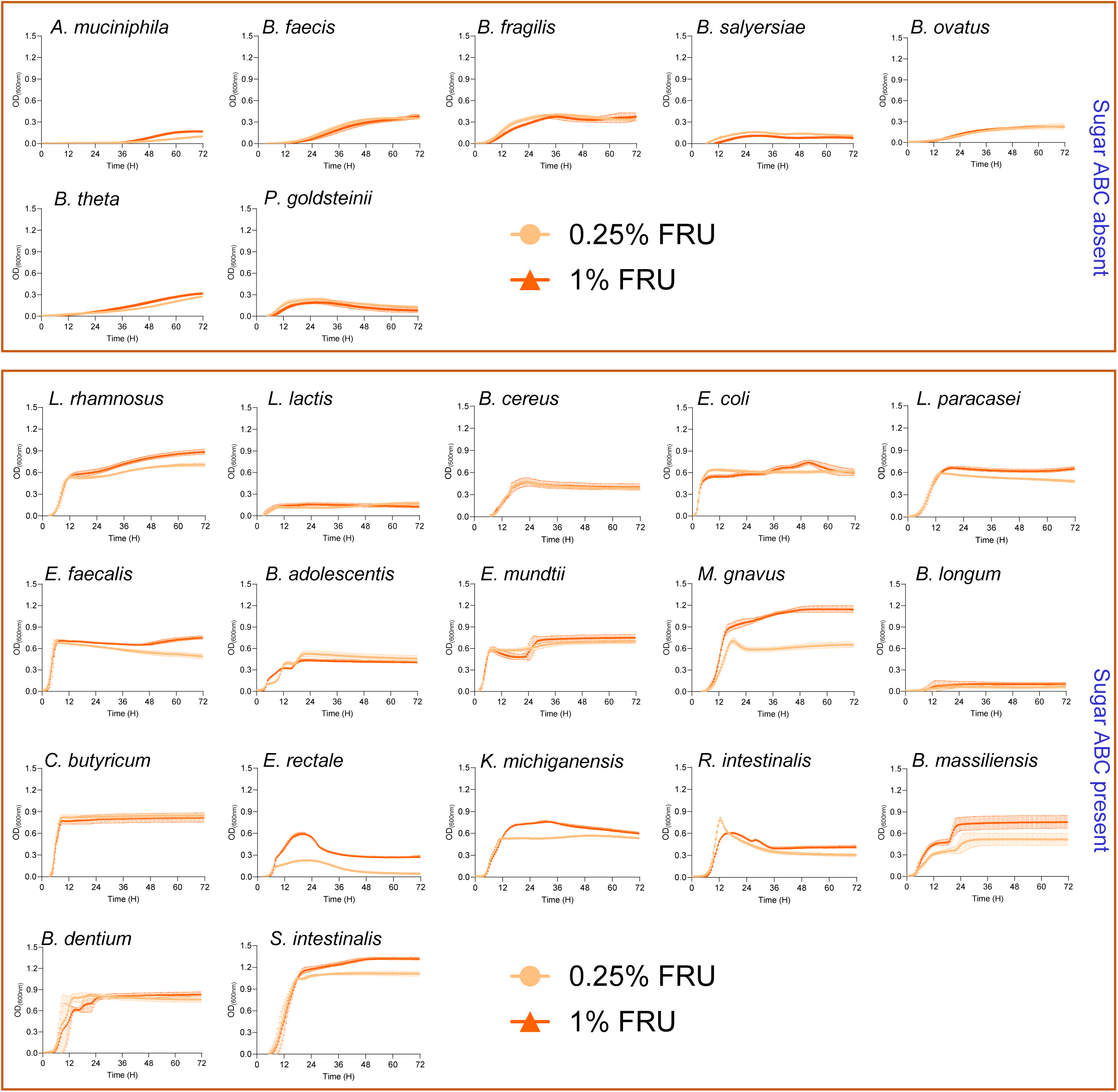
Analysis of the growth (OD_600_) of the indicated strains. Bacteria were cultured in in BHI-NG medium supplemented with 0.25% and 1% fructose (FRU). Graphs represent mean ± SD from two independent experiments (n = 6)

**Supplementary Figure 8.**
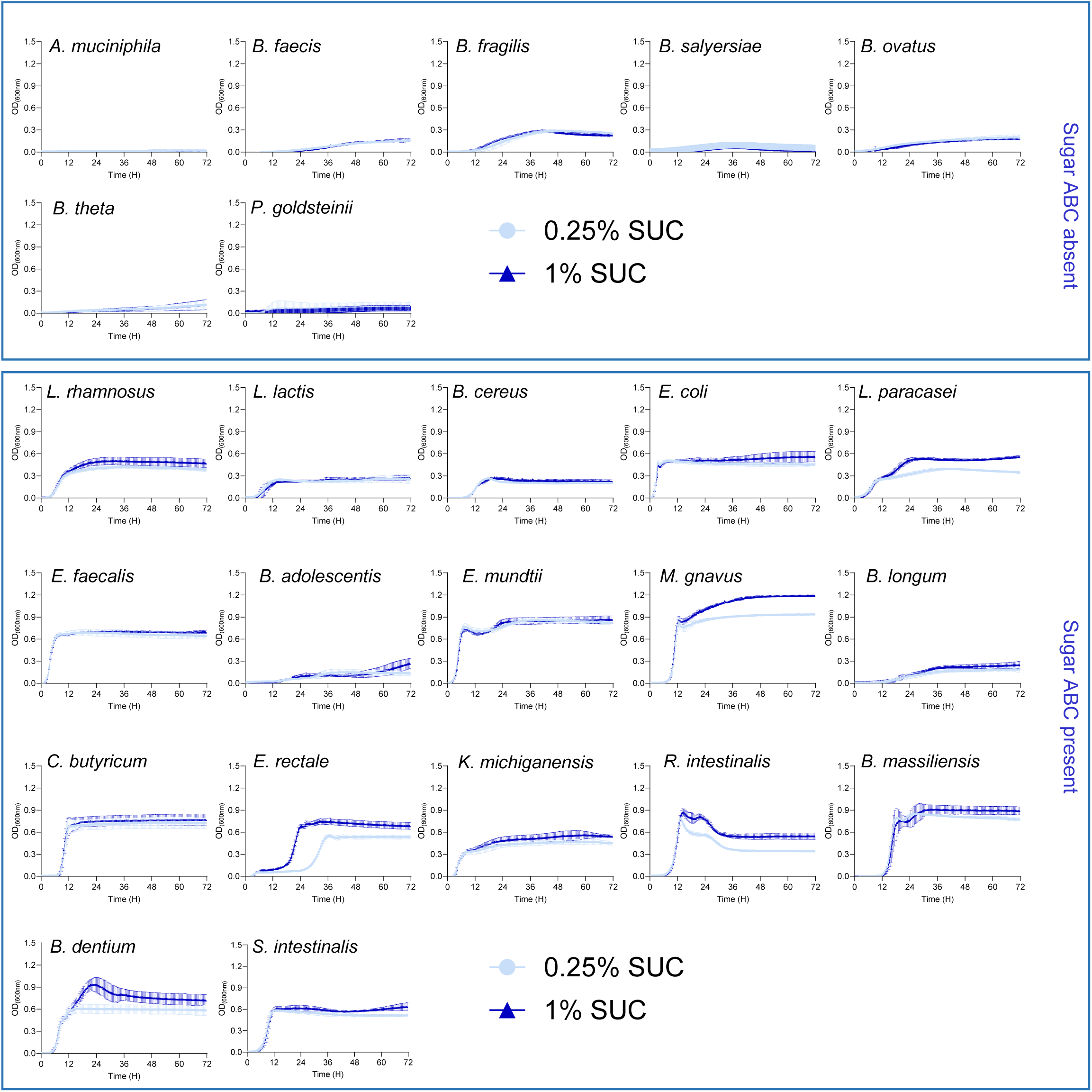
Analysis of the growth (OD_600_) of the indicated strains. Bacteria were cultured in BHI-NG medium supplemented with 0.25% and 1% fructose (SUC). Graphs represent mean ± SD from two independent experiments (n = 6)

**Supplementary Figure 9.**
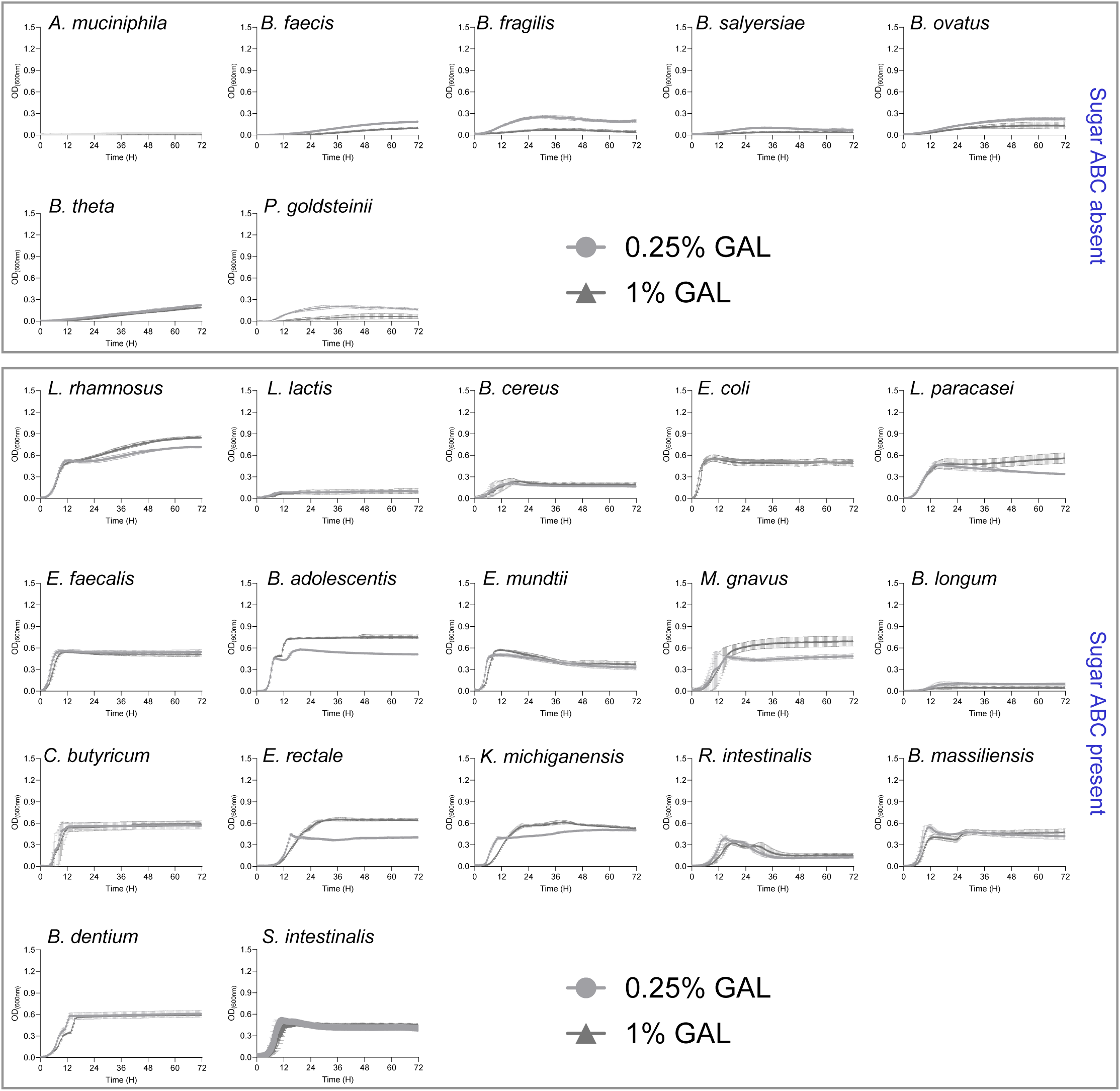
Analysis of the growth (OD_600_) of the indicated strains. Bacteria were cultured in BHI-NG medium supplemented with 0.25% and 1% galactose (GAL). Graphs represent mean ± SD from two independent experiments (n = 6)

**Supplementary Figure 10.**
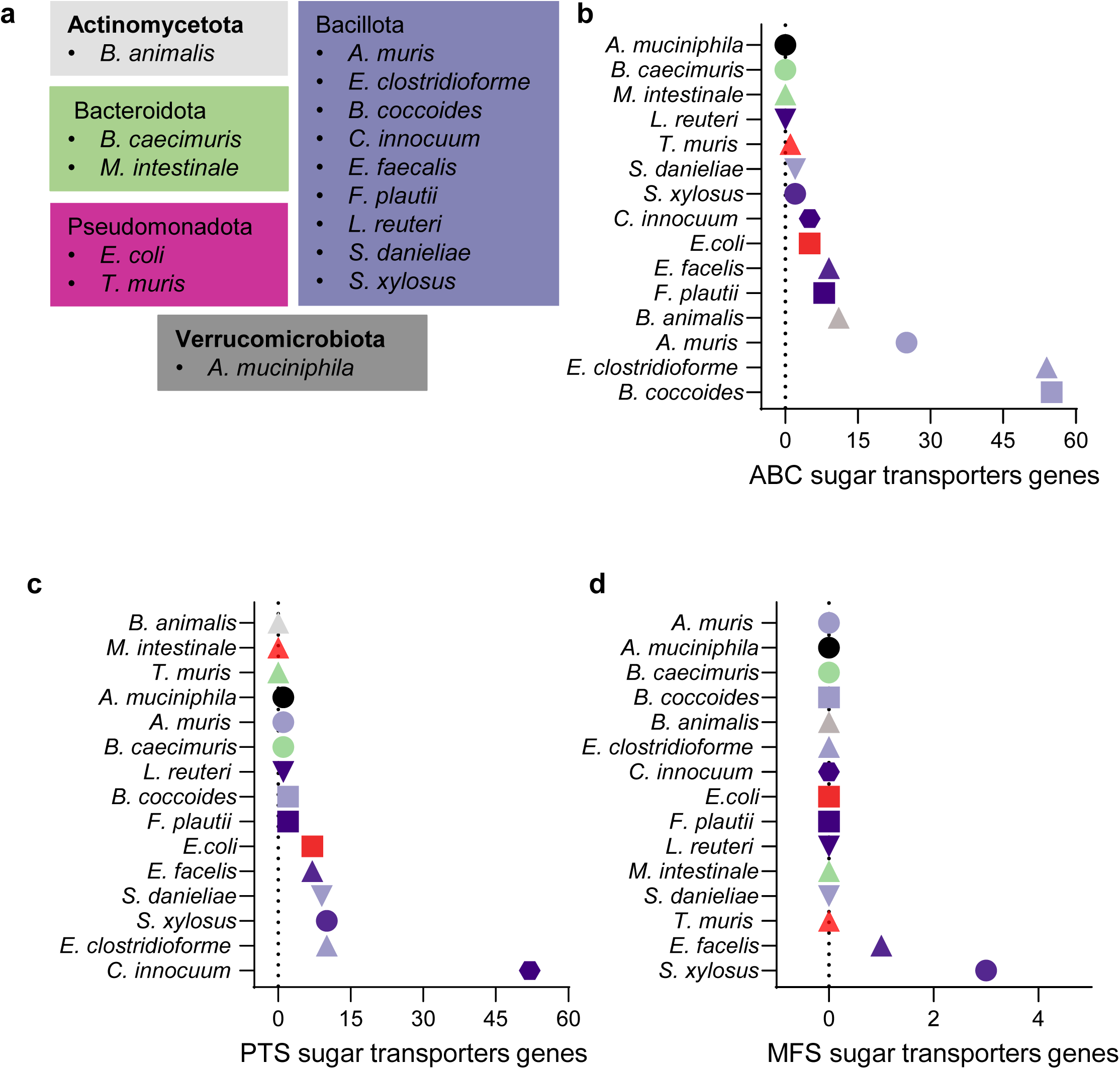
**a**) Table showing the OMM^15^ strains categorized based on their phylum. **b), c), d)** Enumeration of genes involved in sugar ABC, PTS and MFS transporters of the indicated bacteria.

**Supplementary Figure 11.**
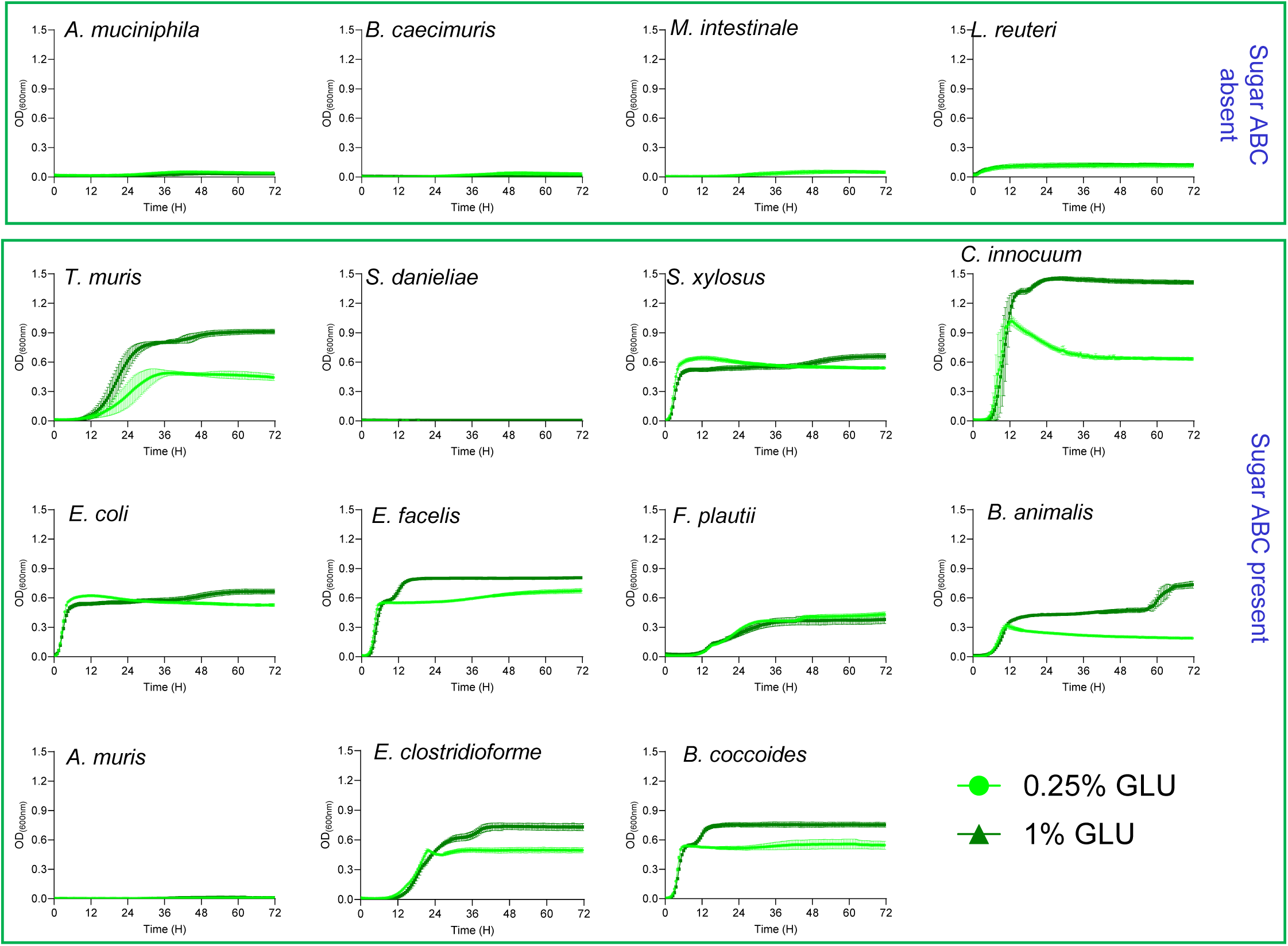
Analysis of the growth (OD_600_) of the indicated OMM^15^ strains. Bacteria were cultured in BHI-NG medium supplemented with 0.25% and 1% glucose. Graphs represent mean ± SD from two independent experiments (n = 6).

**Supplementary Figure 12.**
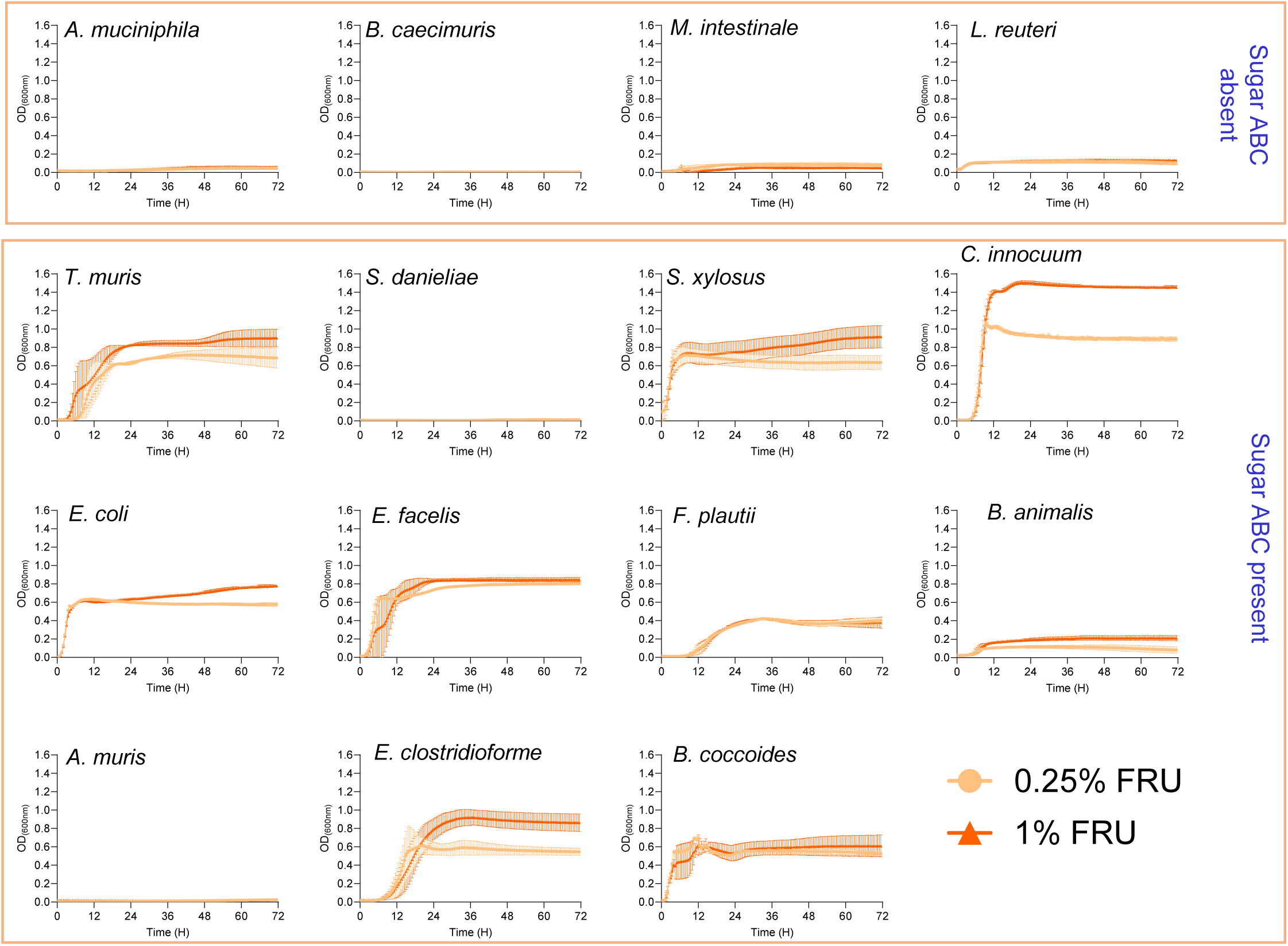
Analysis of the growth (OD_600_) of the indicated OMM^15^ strains. Bacteria were cultured in BHI-NG medium supplemented with 0.25% and 1% fructose. Graphs represent mean ± SD from two independent experiments (n = 6)

**Supplementary Figure 13.**
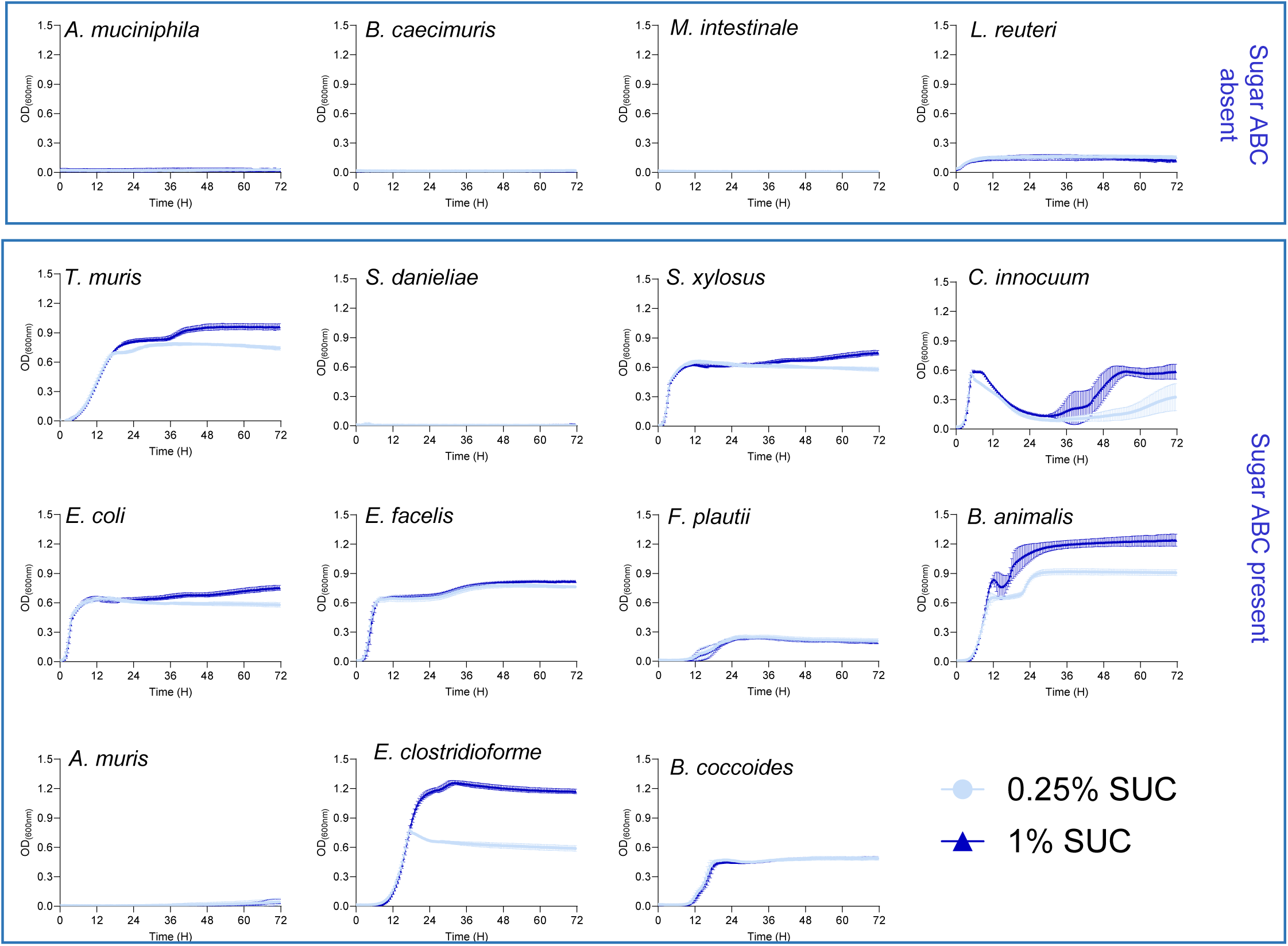
Analysis of the growth (OD_600_) of the indicated OMM^15^ strains. Bacteria were cultured in BHI-NG medium supplemented with 0.25% and 1% fructose. Graphs represent mean ± SD from two independent experiments (n = 6)

**Supplementary Figure 14.**
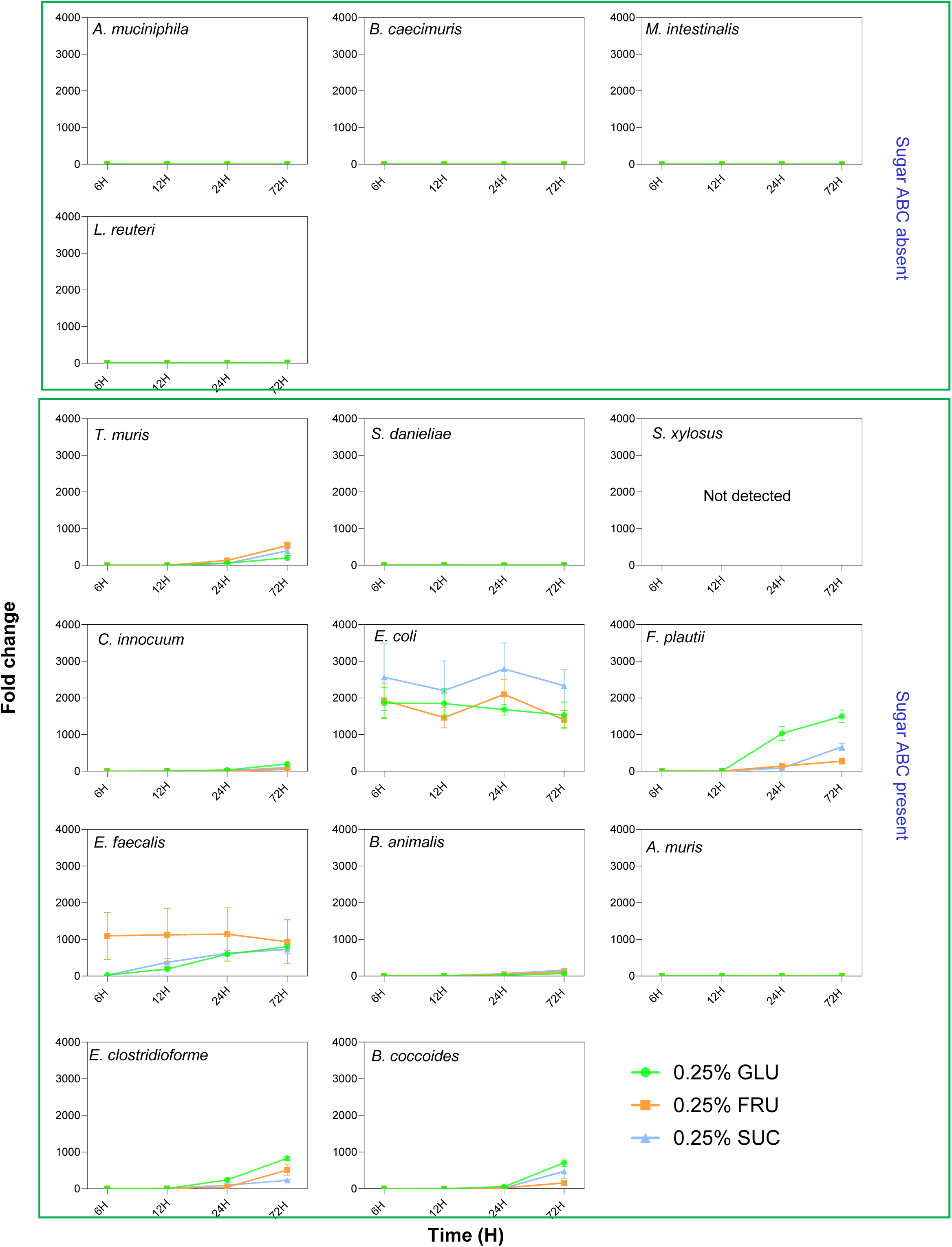
Comparing the fold change in growth for each OMM^15^ species when grown together as an *in vitro* community over a 72-hour period, as determined by quantitative PCR (qPCR). The 0-hour time point (inoculation) was selected as the baseline, and fold change was calculated based on comparisons between 0 to 6, 0 to 12, 0 to 24, and 0 to 72 hours. Bacteria were grown in BHI-NG medium supplemented with 0.25% glucose (GLU), fructose (FRU), or sucrose (SCU). Graphs represent mean ± SD from two independent experiments (n = 6). Data are combined from two independent biological experiments and qPCR was performed in technical duplicate for each species (n = 4).

**Supplementary Figure 15.**
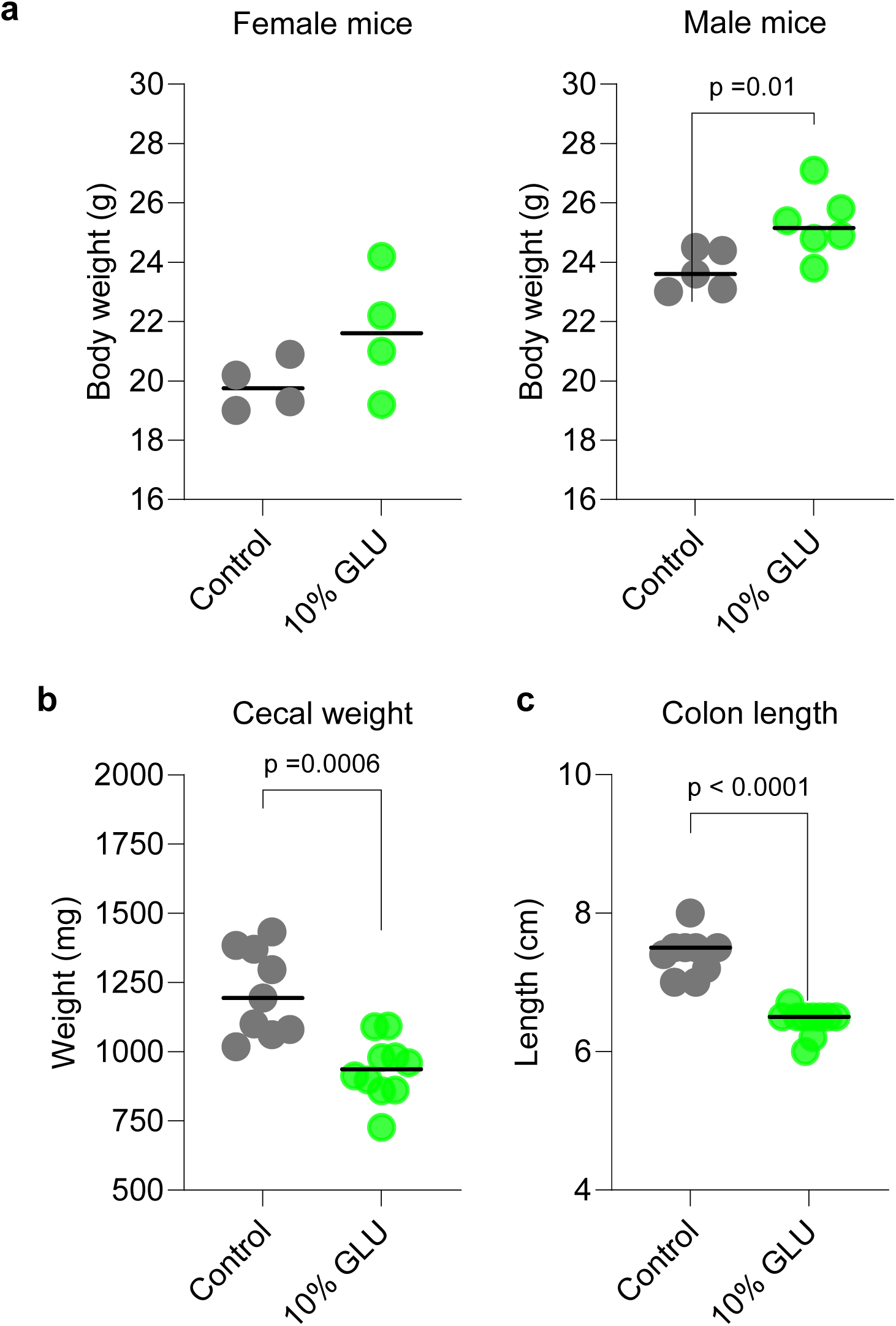
Comparison of **a)** body weight, **b)** cecal weight, and **c)** colon length at day 14 between control mice and mice provided with 10% glucose (GLU) supplemented water. Graphs depict the mean and individual data points for each mouse from one experiment (n = 4 control female mice, n = 4 10% GLU female mice, n = 5 control male mice, and n = 6 10% GLU male mice). Statistical analysis was performed using an unpaired *t*-test with Welch’s correction. *p* < 0.05 was considered statistically significant.

**Supplementary Figure 16.**
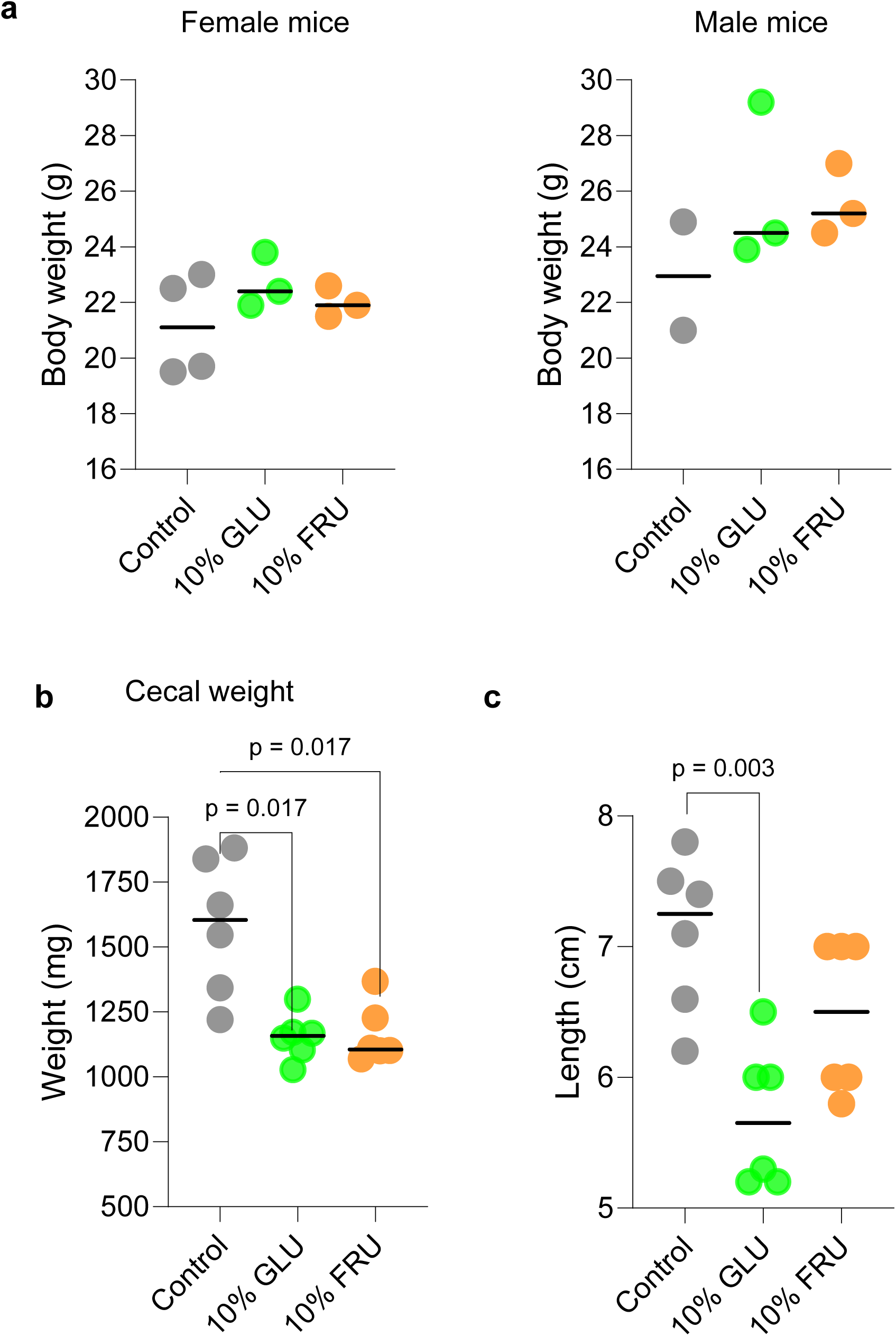
Comparison of **a)** body weight, **b)** cecal weight, and **c)** colon length at day 14 between control mice and mice provided with 10% glucose (GLU) and 10% fructose (FRU) supplemented water. Graphs depict the mean and individual data points for each mouse from one experiment (n = 4 control female mice, n = 3 10% FRU female mice, n = 3 10% GLU female mice n = 2 control male mice, n = 3 10% FRU male mice and n = 3 10% GLU male mice). Statistical analysis was performed using an unpaired *t*-test with Welch’s correction. *p* < 0.05 was considered statistically significant.

**Supplementary Figure 17.**
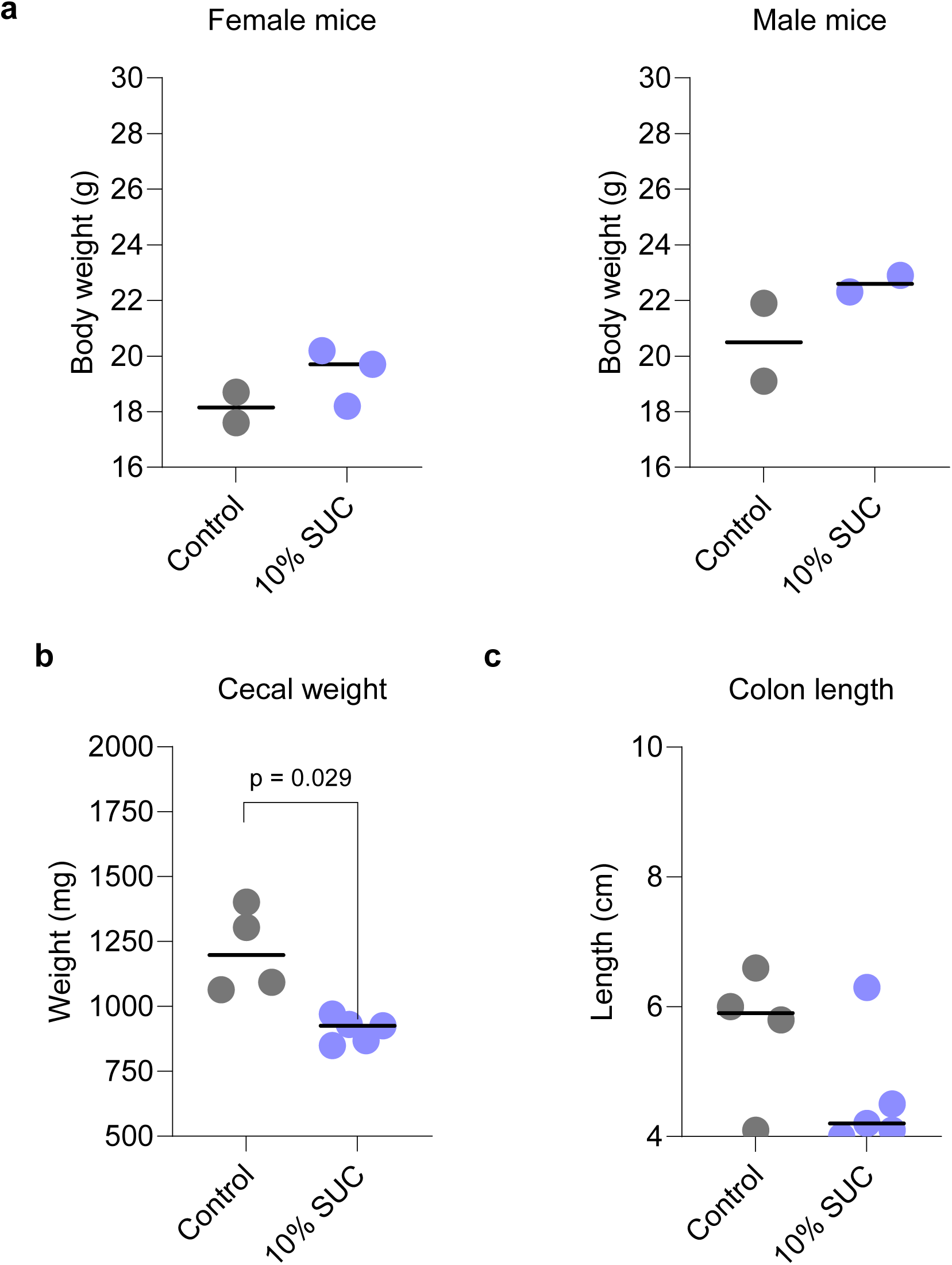
Comparison of **a)** body weight, **b)** cecal weight, and **c)** colon length at day 14 between control mice and mice provided 10% sucrose (SUC) supplemented water. Graphs depict the mean and individual data points for each mouse from one experiment (n = 2 control female mice, n= 3 10% SUC female mice, n = 2 control male mice, n = 2 10% SUC male mice). Statistical analysis was performed using unpaired *t*-test with Welch’s correction. *p* < 0.05 was considered statistically significant.

**Supplementary Figure 18.**
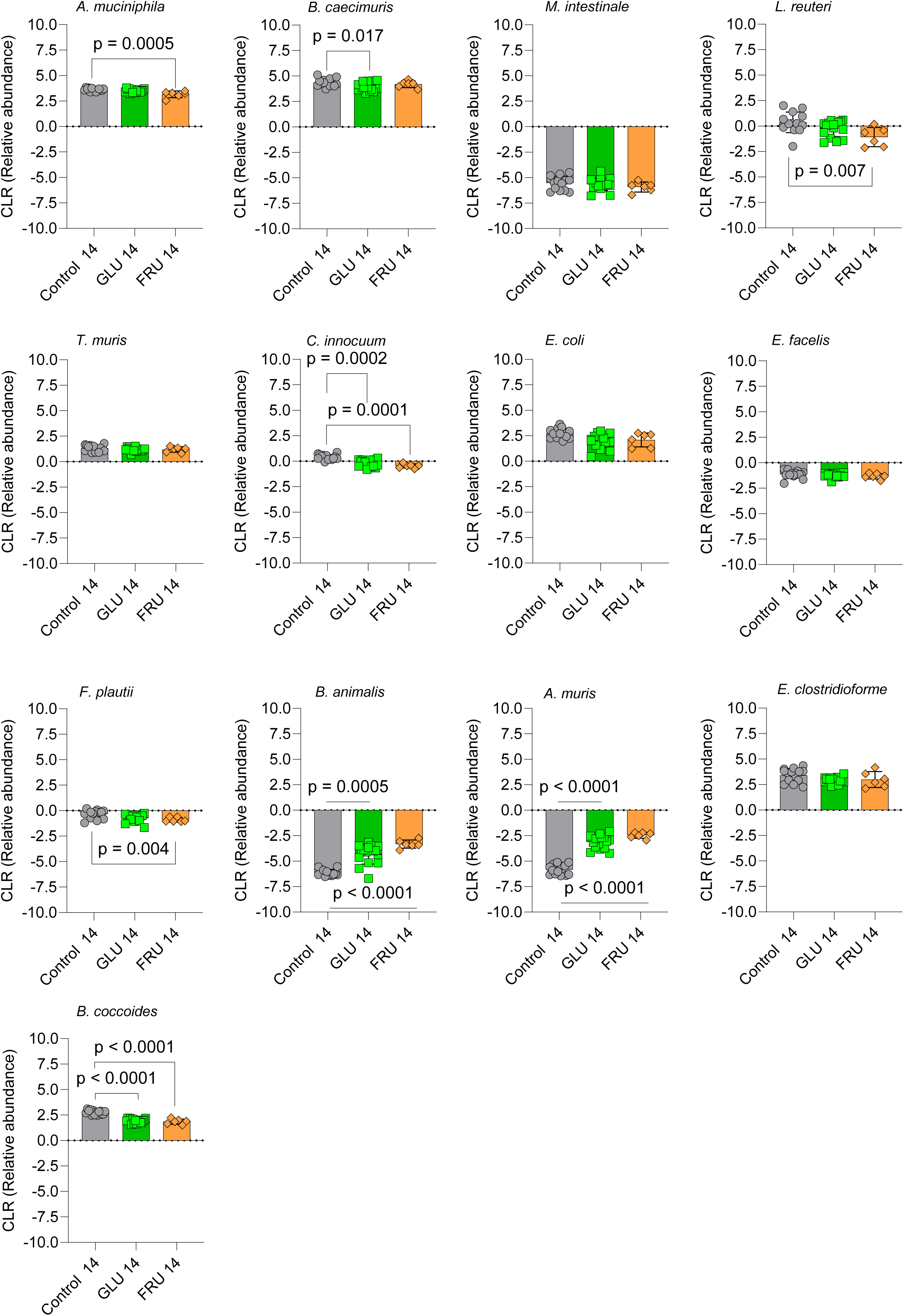
Comparison of the CLR-transformed relative abundance of OMM^15^ community members on day 14 between control mice and those provided with 10% GLU or FRU supplemented water. Data are combined from two independent experiments (*n* = 8 control female mice, *n* = 7 control male mice, *n* = 7 GLU female mice, *n* = 9 GLU male mice, *n* = 3 FRU female mice, *n* = 3 FRU male mice). Statistical analysis was performed using the Kruskal–Wallis test with Dunn’s multiple comparisons, with adjusted p values reported. *S. danieliae* and *S. xylosus* were undetectable by 16S rRNA sequencing. *P* < 0.05 was considered statistically significant.

**Supplementary Figure 19.**
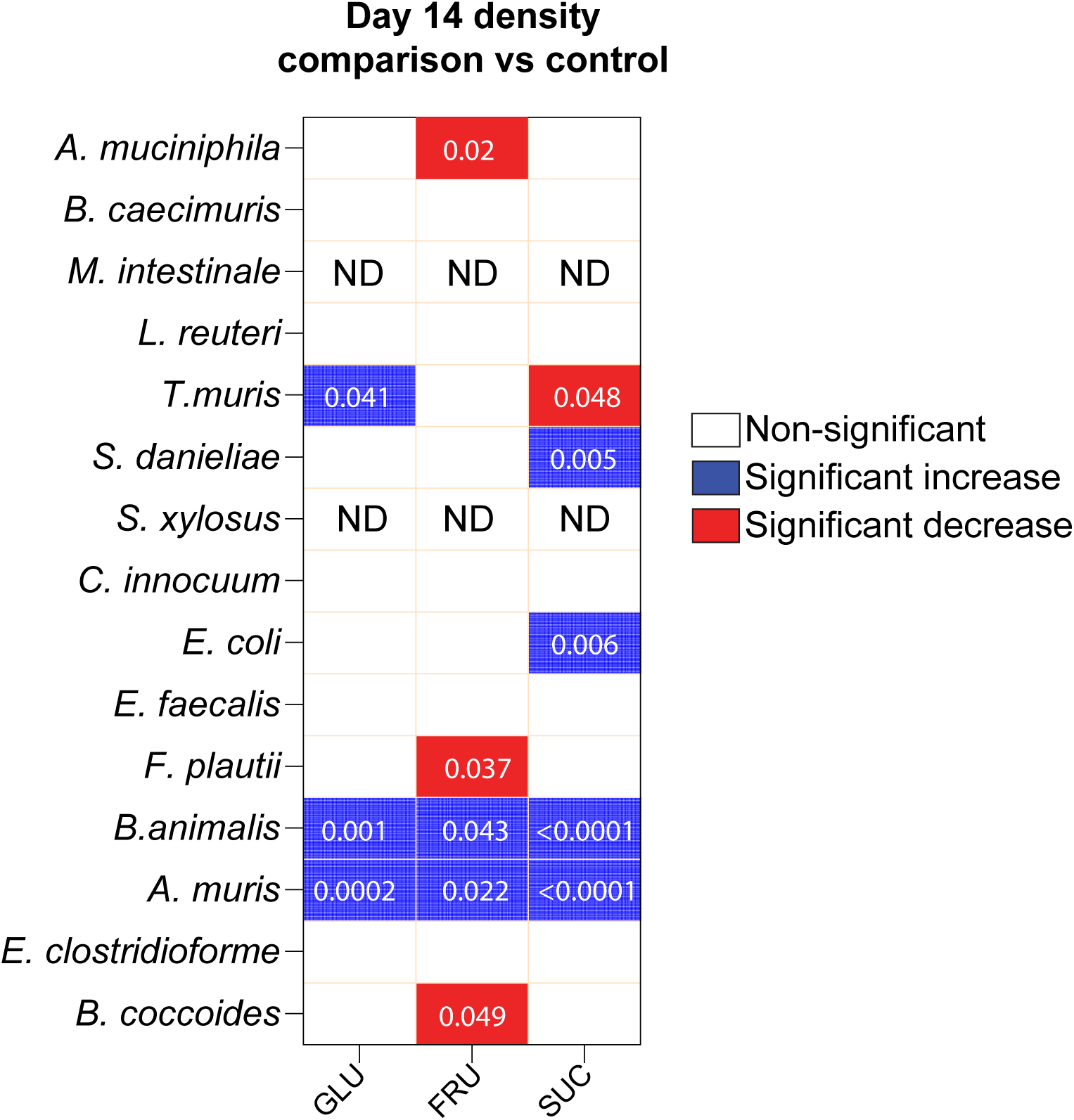
Differences in density of OMM^15^ species using normalized cT values (cT / fecal weight) from qPCR data of OMM^15^ community members on day 14 between control mice and those provided with 10% glucose (GLU), fructose (FRU), or sucrose (SUC) supplemented water. Data were combined from three independent experiments (control: n = 10 females, n = 9 males; GLU: n = 7 females, n = 9 males; FRU: n = 3 females, n = 3 males; SUC: n = 3 females, n = 2 males). ND, not detected, *M. intestinale* and *S. xylosus* were below the qPCR detection limit. Statistical analysis was performed using the Kruskal–Wallis test with Dunn’s multiple comparisons. Directionality of differences with adjusted p-values < 0.05 is indicated by color.

**Supplementary Figure 20.**
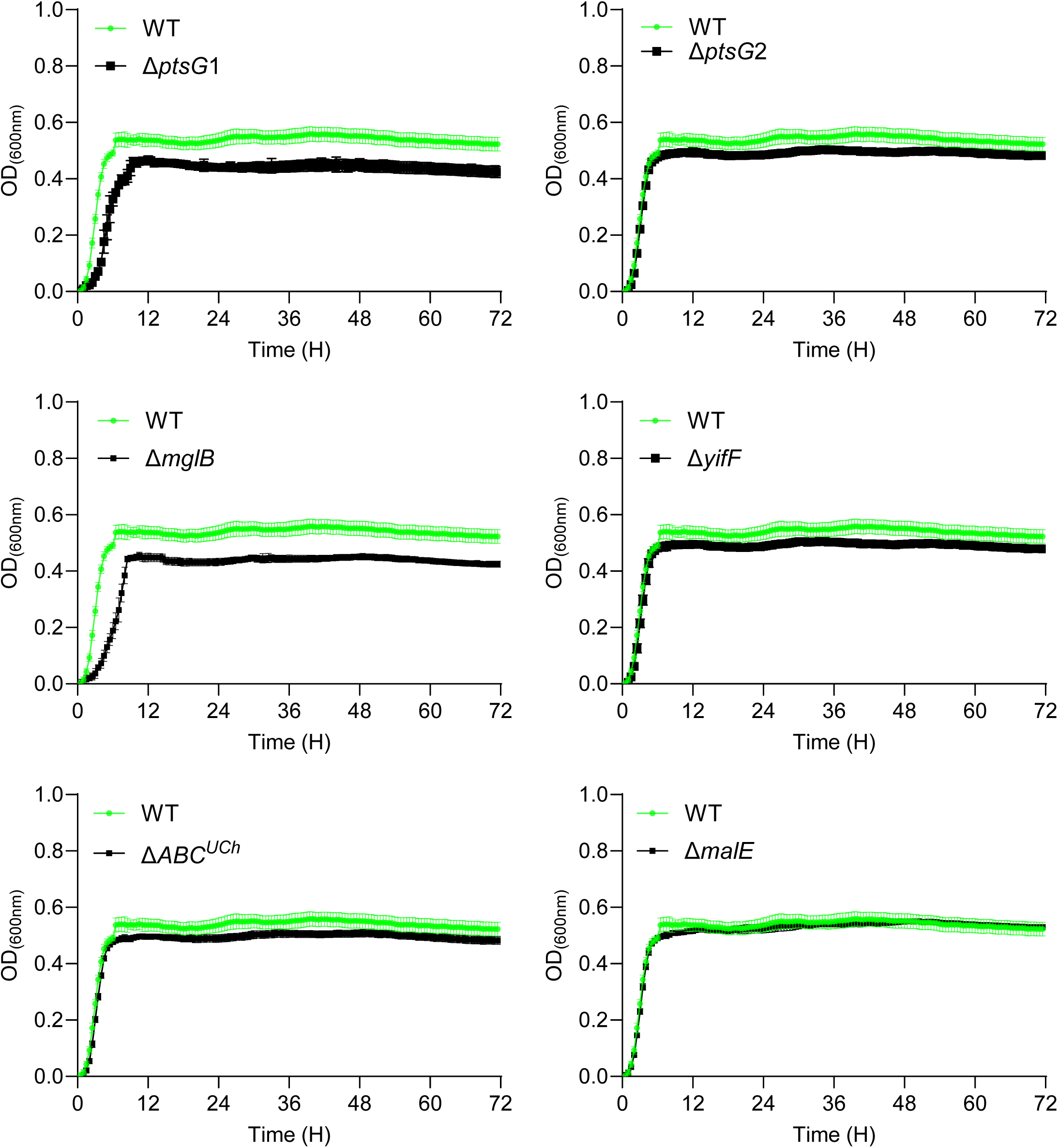
Analysis of the growth (OD_600_) of WT *E. coli* MT1B1 and its indicated sugar ABC and PTS transporters permease mutants. Bacteria were cultured in BHI-NG medium supplemented with 0.25% glucose. Graphs represent mean ± SD from two independent experiments (n = 6).

**Supplementary Figure 21.**
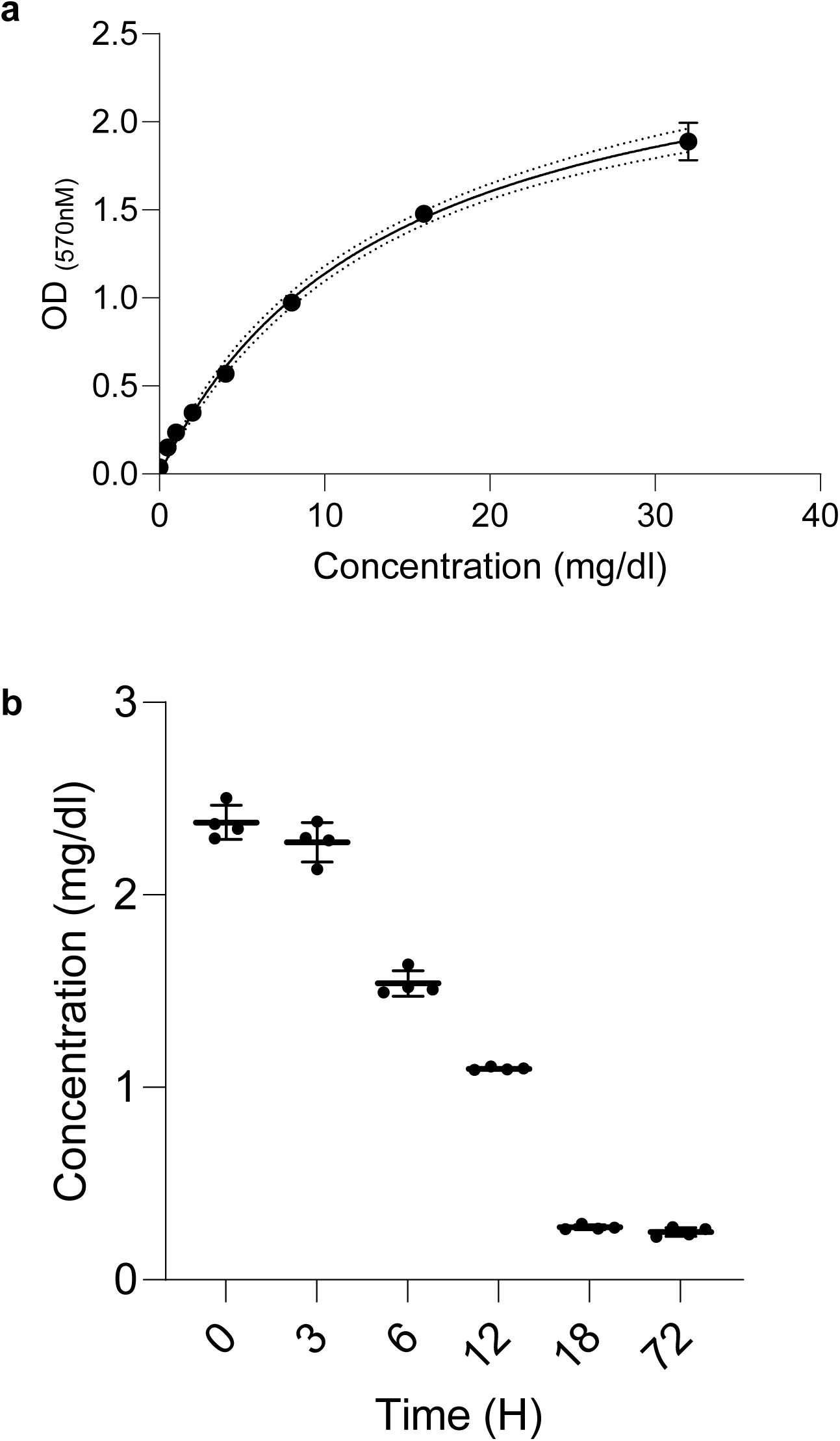
**a)** Standard curve generated using the Invitrogen glucose colorimetric detection kit. Standards were used in in duplicates (n=2). **b)** Graph showing glucose concentrations over a period of 72-hour in an OMM^15^ community cultured in 0.25%glucose (GLU). Samples were diluted 100 folds. Graphs represent mean ± SD from two independent experiments (n = 6)

**Supplementary table 1:**
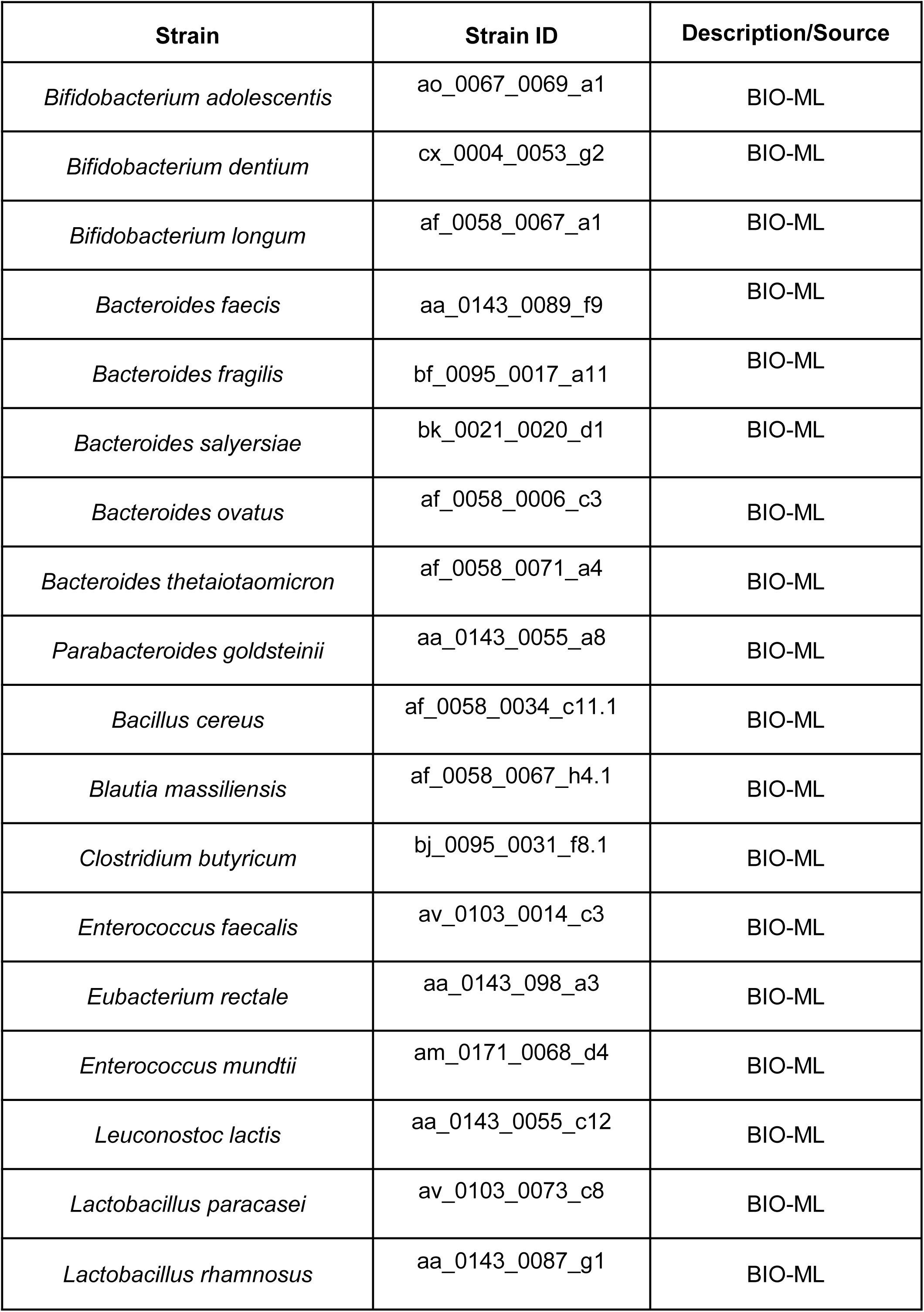
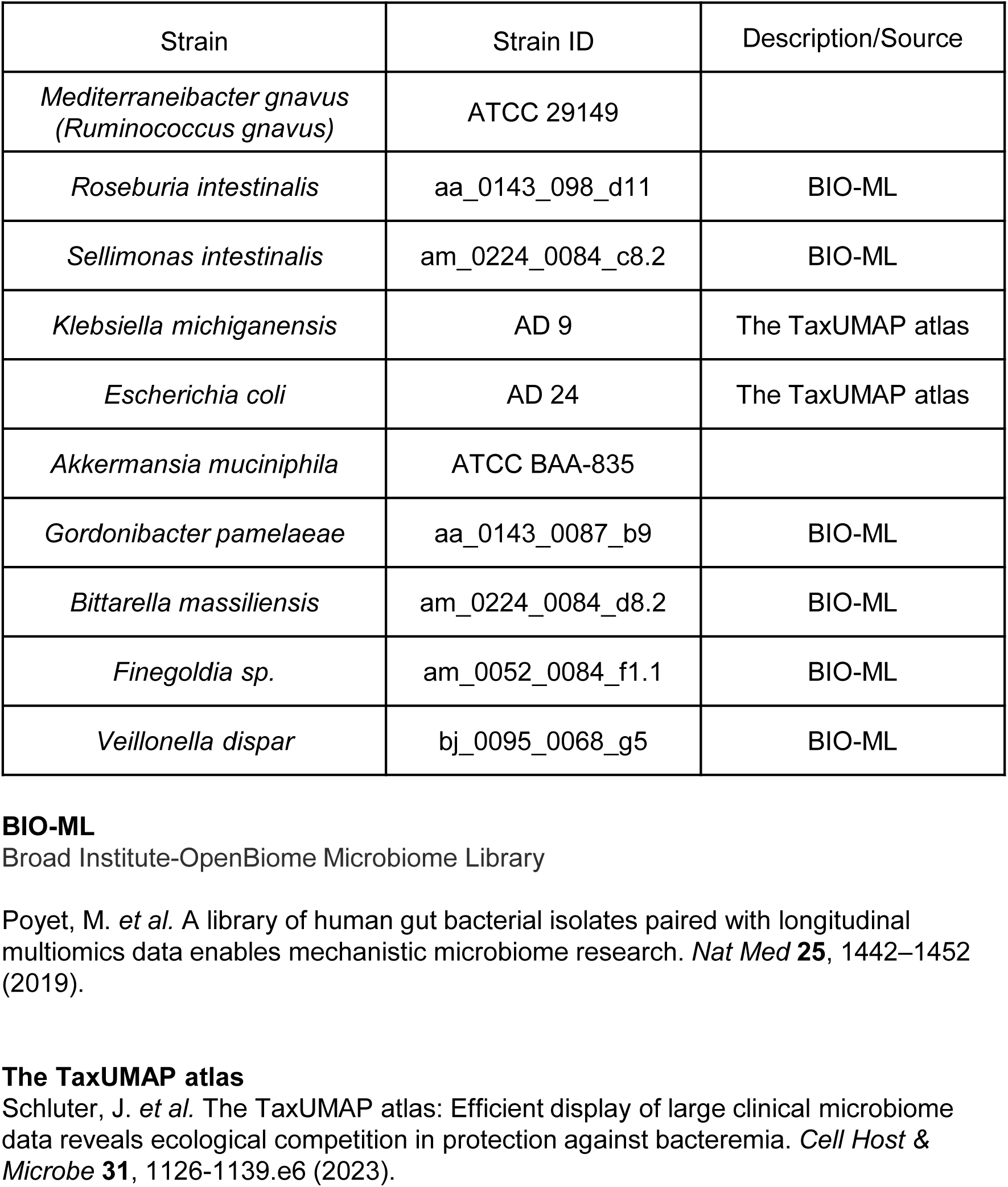
List of bacterial species.

| Strain | Strain ID | Description/Source |
| --- | --- | --- |
| *Bifidobacterium adolescentis* | ao_0067_0069_a1 | BIO-ML |
| *Bifidobacterium dentium* | cx_0004_0053_g2 | BIO-ML |
| *Bifidobacterium longum* | af_0058_0067_a1 | BIO-ML |
| *Bacteroides faecis* | aa_0143_0089_f9 | BIO-ML |
| *Bacteroides fragilis* | bf_0095_0017_a11 | BIO-ML |
| *Bacteroides salyersiae* | bk_0021_0020_d1 | BIO-ML |
| *Bacteroides ovatus* | af_0058_0006_c3 | BIO-ML |
| *Bacteroides thetaiotaomicron* | af_0058_0071_a4 | BIO-ML |
| *Parabacteroides goldsteinii* | aa_0143_0055_a8 | BIO-ML |
| *Bacillus cereus* | af_0058_0034_c11.1 | BIO-ML |
| *Blautia massiliensis* | af_0058_0067_h4.1 | BIO-ML |
| *Clostridium butyricum* | bj_0095_0031_f8.1 | BIO-ML |
| *Enterococcus faecalis* | av_0103_0014_c3 | BIO-ML |
| *Eubacterium rectale* | aa_0143_098_a3 | BIO-ML |
| *Enterococcus mundtii* | am_0171_0068_d4 | BIO-ML |
| *Leuconostoc lactis* | aa_0143_0055_c12 | BIO-ML |
| *Lactobacillus paracasei* | av_0103_0073_c8 | BIO-ML |
| *Lactobacillus rhamnosus* | aa_0143_0087_g1 | BIO-ML |
| *Mediterraneibacter gnavus*<br>(*Ruminococcus gnavus*) | ATCC 29149 |  |
| *Roseburia intestinalis* | aa_0143_098_d11 | BIO-ML |
| *Sellimonas intestinalis* | am_0224_0084_c8.2 | BIO-ML |
| *Klebsiella michiganensis* | AD 9 | The TaxUMAP atlas |
| *Escherichia coli* | AD 24 | The TaxUMAP atlas |
| *Akkermansia muciniphila* | ATCC BAA-835 |  |
| *Gordonibacter pamelaeae* | aa_0143_0087_b9 | BIO-ML |
| *Bittarella massiliensis* | am_0224_0084_d8.2 | BIO-ML |
| *Finegoldia sp.* | am_0052_0084_f1.1 | BIO-ML |
| *Veillonella dispar* | bj_0095_0068_g5 | BIO-ML |

**Supplementary table 2:** Glycolytic genes and their function.

| Gene | Pathway | Function | Reference |
| --- | --- | --- | --- |
| Glyceraldehyde 3-phosphate dehydrogenase<br>**GAPDH** | EMP | Conversion of glyceraldehyde-3-phosphate to 1,3-bisphosphoglycerate | 1,2 |
| Hexokinase (ROK protein family)<br>**HK** | EMP, PPP, ED | catalyzes the phosphorylation of glucose by ATP to glucose-6-phosphate | 1,2 |
| Pyruvate kinase<br>**PK** | EMP | conversion of phosphoenolpyruvate to pyruvate | 1,2 |
| Phosphoglucomutase<br>**PGM** | EMP, PPP, ED | PGM converts Glucose-1-Phosphate released from glycogen into Glucose-6-Phosphate, which can then be used in the glycolytic pathways | 1,2,3,4 |
| Phosphofructokinase<br>**PFK** | EMP | conversion of fructose-6-phosphate to fructose-1,6-diphosphate | 1,2 |
| Ribose-5-phosphate isomerase<br>**RPI** | PPP | conversion of ribose-5-phosphate to ribulose-5-phosphate | 1,4 |
| Transketolase<br>**TK** | EMP, PPP | Linking EMP to PPP pathway | 1,4 |
| Phosphogluconate dehydratase<br>**6PGDH** | ED | 6-phosphogluconate (6-PG) is dehydrated to 2-keto-3-deoxy-6-phosphogluconate | 1,3 |
| 2-dehydro-3-deoxy-phosphogluconate aldolase<br>**KDPG** | ED | Conversion of 2-Keto-3-deoxy-6-phosphogluconate (KDPG) to pyruvate and D-glyceraldehyde-3-phosphate | 1,3 |

1.Wolfe, A. J. Glycolysis for the Microbiome Generation. *Microbiol Spectr* **3**, 10.1128/microbiolspec.MBP-0014–2014 (2015).
2.Sánchez-Pascuala, A., de Lorenzo, V. & Nikel, P. I. Refactoring the Embden–Meyerhof–Parnas Pathway as a Whole of Portable GlucoBricks for Implantation of Glycolytic Modules in Gram-Negative Bacteria. *ACS Synth. Biol.* **6**, 793–805 (2017).
3.Peekhaus, N. & Conway, T. What's for Dinner?: Entner-Doudoroff Metabolism in Escherichia coli. *Journal of Bacteriology* **180**, 3495 (1998).
4.Rashida, Z. & Laxman, S. The pentose phosphate pathway and organization of metabolic networks enabling growth programs. *Current Opinion in Systems Biology* **28**, 100390 (2021).

**Supplementary table 3:** Sugar - sweetened beverages consumed by HCT patients.

| **FNDDS Food Code** | **Description** | **Prevalence in 1009 samples** | **Sugars per g (g)** |
| --- | --- | --- | --- |
| 92410610 | Soft drink, ginger ale | 206 | 1.01 |
| 95320200 | Sports drink (Gatorade G) | 142 | 0.79 |
| 92410310 | Soft drink, cola | 93 | 0.93 |
| 92309000 | Tea, iced, bottled, black | 88 | 0.86 |
| 95103000 | Nutritional drink or shake, ready-to-drink (Ensure) | 83 | 0.38 |
| 92410510 | Soft drink, fruit flavored, caffeine free | 50 | 0.88 |
| 92513000 | Fruit flavored smoothie drink, frozen, no dairy | 46 | 0.86 |
| 95103010 | Nutritional drink or shake, ready-to-drink (Ensure Plus) | 35 | 0.18 |
| 95104000 | Nutritional drink or shake, ready-to-drink, sugar free (Glucerna) | 8 | 0.13 |
| 95101010 | Nutritional drink or shake, ready-to-drink (Boost Plus) | 6 | 0.23 |
| 95322200 | Sports drink, low calorie (Gatorade G2) | 5 | 0.63 |
| 95220000 | Nutritional powder mix, NFS | 5 | 0.24 |
| 92510610 | Fruit juice drink | 3 | 1.00 |
| 92530610 | Fruit juice drink, with high vitamin C | 3 | 0.90 |
| 92101900 | Coffee, Latte | 3 | 0.45 |
| 92510955 | Lemonade, fruit juice drink | 2 | 0.96 |
| 92306800 | Tea, hot, chai, with milk | 1 | 0.74 |
| 95330100 | Fluid replacement, electrolyte solution | 1 | 0.69 |
| 95106000 | Nutritional drink or shake, ready-to-drink (Muscle Milk) | 1 | 0.07 |
| 95202000 | Nutritional powder mix (Muscle Milk) | 1 | 0.07 |

Sugar-sweetened beverages were defined as beverages with added sugar, not including enteral formulae. Sugars per gram of food are reported, as they appear in the FNDDS database.

**Supplementary table 4:** MM^15^ strains used in the study.

| Strain | Strain ID |
| --- | --- |
| *Acutalibacter muris* | MM12 KB18 (DSM 26090) |
| *Akkermansia muciniphila* | MM12 YL44 (DSM 26127) |
| *Bacteroides caecimuris* | MM12 I48 (DSM 26085) |
| *Bifidobacterium animalis subsp. animalis* | MM12 YL2 (DSM 26074) |
| *Blautia coccoides* | MM12 YL58 (DSM 26115) |
| *Clostridium innocuum* | MM12 I46 (DSM 26113) |
| *Enterocloster clostridioforme* | MM12 YL32 (DSM 26114) |
| *Enterococcus faecalis* | MM12 KB1 (DSM 32036) |
| *Escherichia coli* | Mt1B1 (DSM 28618) |
| *Flavonifractor plautii* | MM12 YL31 (DSM 26117) |
| *Limosilactobacillus reuteri* | MM12 I49 (DSM 32035) |
| *Muribaculum intestinale* | MM12 YL27 (DSM 28989) |
| *Staphylococcus xylosus* | 33-ERD13C (DSM 28566) |
| *Streptococcus danieliae* | ERD01G (DSM 22233) |
| *Turicimonas muris* | MM12 YL45 (DSM 26109) |

**Supplementary table 5:** List of *Escherichia coli* MT1B1 mutants.

| Mutant | Description | Reference |
| --- | --- | --- |
| *ΔptsG1 (EIIC1)* | Permease of first PTS transporter responsible for the transport of sugar molecules. Cm<sup>r</sup> | 1 |
| *ΔptsG2 (EIIC2)* | Permease of second PTS transporter responsible for the transport of sugar molecules. Cm<sup>r</sup> | 1 |
| *ΔmglB* | *mglB* is a high affinity uptake system of the MglBAC ABC transporter system, which is responsible for the uptake of D-galactose, glucose and methyl-D-galactoside. Cm<sup>r</sup> | 2 |
| *ΔABC<sup>Uch</sup>* | Unknown sugar ABC permease. Cm<sup>r</sup> | This study |
| *ΔyjfF* | *yjfF* is a permease of YtfQRT-YjfF ABC transporter system which is responsible for the uptake of Galactofuranose. Cm<sup>r</sup> | 3 |
| *ΔmalE* | *malE* is a high affinity uptake system of the MalEFGK ABC transporter system which is responsible for the uptake of maltose and maltodextrin. Cm<sup>r</sup> | 3 |

This strain possesses six putative genes encoding components of active sugar transporters, including two PTS permeases (EIIC1 and EIIC2), both annotated as *ptsG* in our screen but referred to here as *ptsG1* and *ptsG2* for clarity. Additionally, it contains four sugar ABC transporter components: two high affinity transport systems *mglB* and *malE*; *yjfF*, an inner membrane sugar ABC protein; and one uncharacterized gene functionally assigned as a sugar ABC permease (*ABC<sup>UCh</sup>*).
1.McCoy, J. G., Levin, E. J. & Zhou, M. Structural insight into the PTS sugar transporter EIIC. *Biochimica et biophysica acta* **1850**, 577 (2014).
2.Ferenci, T. Adaptation to life at micromolar nutrient levels: the regulation of *Escherichia coli* glucose transport by endoinduction and cAMP. *FEMS Microbiology Reviews* **18**, 301–317 (1996).
3.Blattner, F. R. *et al.* The complete genome sequence of *Escherichia coli* K-12. *Science* **277**, 1453–1462 (1997).

**Supplementary table 6:** List of plasmids used to generate *Escherichia coli* MT1B1 mutants.

| Plasmid | Resistance | Reference |
| --- | --- | --- |
| pkd46 | Amp<sup>r</sup> | 1 |
| pkd3 | Amp<sup>r</sup>, Cm<sup>r</sup> | 1 |

Amp<sup>r</sup>– tetracycline resistance, Cm<sup>r</sup> – chloramphenicol resistance
1.One-step inactivation of chromosomal genes in *Escherichia coli* K-12 using PCR products | PNAS.

**Supplementary table 7:**
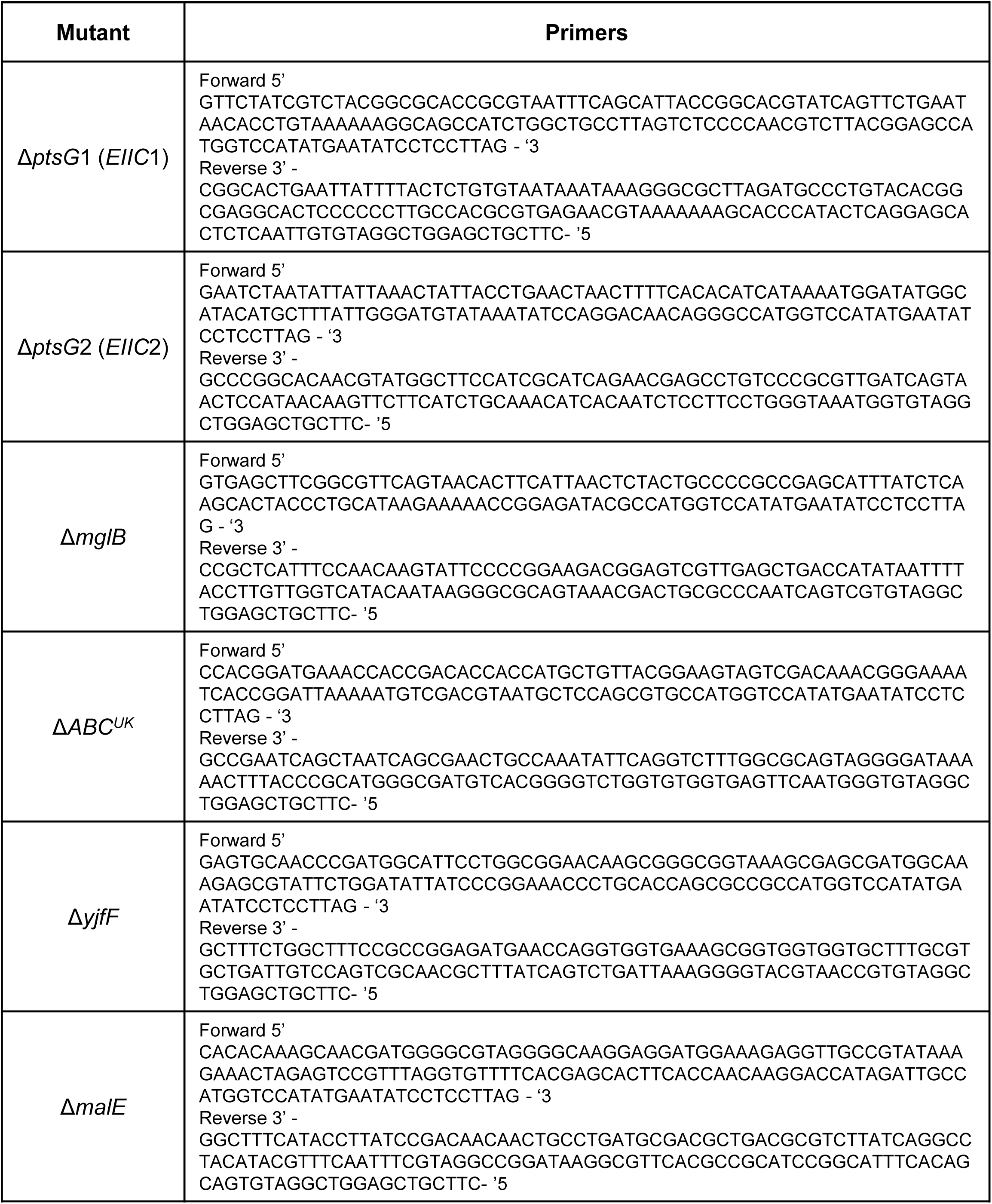
List of primers used to generate *Escherichia coli* MT1B1 mutants.

**Supplementary table 8:**
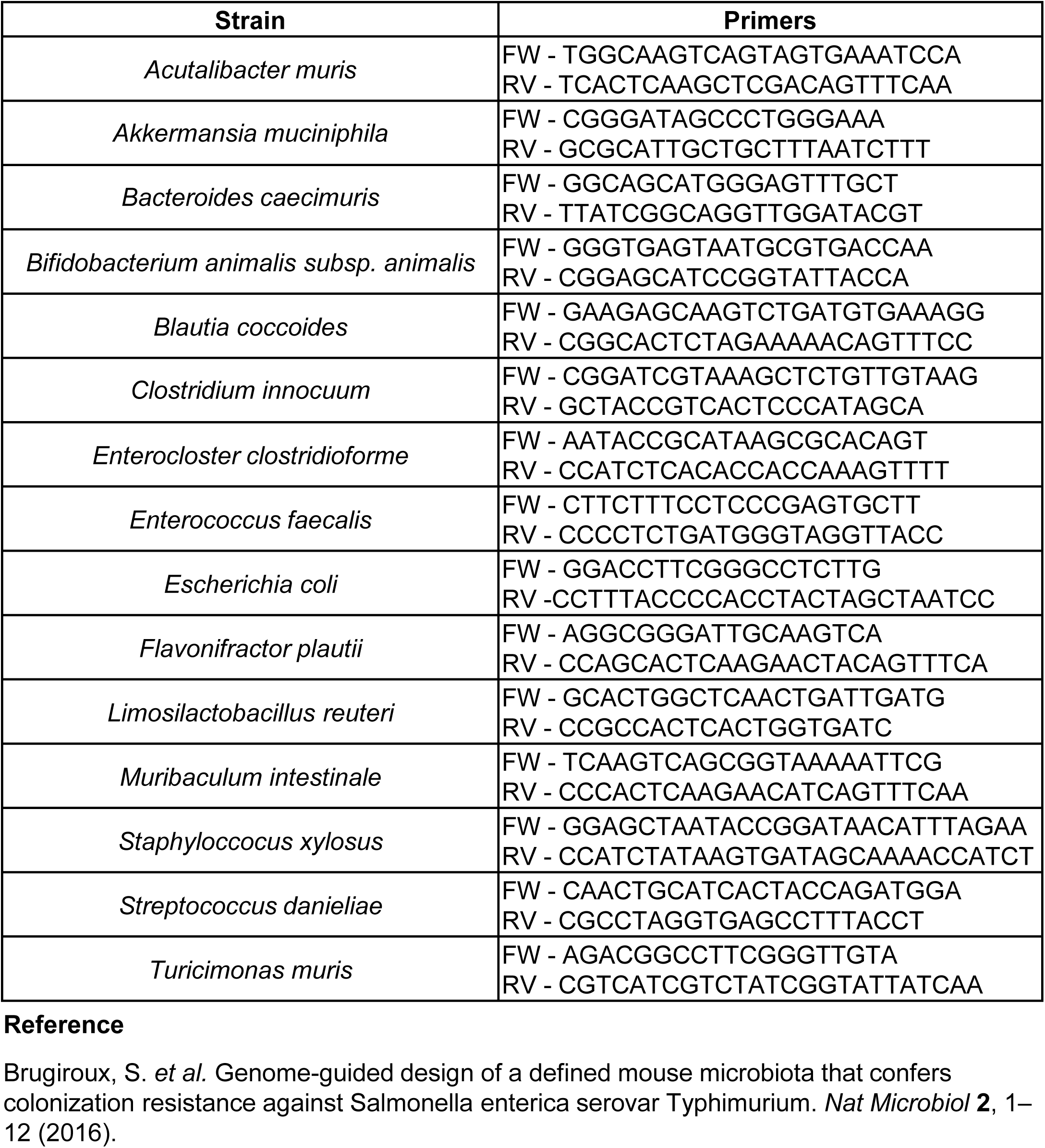
List of MM^15^ qPCR primers used in this study.

